# Diversification of Polynucleotide Kinase Clp1 Family Proteins and Their Possible Origins in Eukaryotic Evolution

**DOI:** 10.1101/2023.03.12.532321

**Authors:** Motofumi Saito, Rerina Inose, Asako Sato, Masaru Tomita, Akio Kanai

## Abstract

The Clp1 family proteins, consisting of the Clp1 and Nol9/Grc3 groups, have polynucleotide kinase (PNK) activity at the 5′ end of RNA strands and are important enzymes in the processing of some precursor RNAs. However, it remains unclear how this enzyme family diversified in the eukaryotes. We performed a large-scale molecular evolutionary analysis of the full-length genomes of 358 eukaryotic species and identified a group of protist proteins that we consider to be the evolutionary origin of Clp1 family proteins. First, we detected an average of 4.2 Clp1 family genes in members of in the protist phylum Euglenozoa, arising from gene duplication. For example, in *Trypanosoma brucei*, there are three genes of the Clp1 group and one gene of the Nol9/Grc3 group. In the Clp1 group proteins encoded by these three genes, the C-terminal domains have been replaced by unique characteristics domains, so we designated these proteins *Tb*-Clp1-t1, *Tb*-Clp1-t2, and *Tb*-Clp1-t3. Experimental validation showed that only *Tb*-Clp1-t2 has PNK activity against RNA strands. In a phylogenetic analysis of the PNK domain, *Tb*-Clp1-t2 mapped to the earliest position on the tree. Interestingly, *Tb*-Clp1-t1, for which no PNK activity was detected, was located on the phylogenetic tree near fungal *Saccharomyces cerevisiae* Clp1, which also lacks PNK activity. The analysis also revealed that in the higher eukaryotes, such as humans and plants, the Clp1 proteins diversified with the creation of isoforms by alternative splicing.

## Introduction

Cleavage factor polyribonucleotide kinase subunit 1 (Clp1) is an enzyme with polynucleotide kinase (PNK) activity, which transfers the phosphate group of ATP to the 5′ end of RNA (Weitzer and Martinez 2007). This enzyme is mainly involved in splicing precursor transfer RNA (pre-tRNA) (Weitzer and Martinez 2007) and the formation of the 3′-end of messenger RNA (mRNA) (Minvielle-Sebastia et al. 1997; de Vries et al. 2000). Many eukaryotic and archaeal tRNA genes contain intron(s), and precise pre-tRNA splicing reactions are required to produce functional mature tRNAs (Abelson et al. 1998). For example, consider a pre-tRNA gene with one intron in its anticodon loop region. The removal of the intron is catalysed by the tRNA splicing endonuclease, and the ligation of the resulting two exons is either via the phosphate group at the 3′ end of the 5′ tRNA exon (the 3′-phosphate ligation pathway) or via the addition of a new phosphate group to the 5′ end of the 3′ tRNA exon (the 5′-phosphate ligation pathway) (Popow et al. 2012). Clp1 is the enzyme that adds a phosphate group to the 5′ end of the tRNA 3′ exon in the 5′-phosphate ligation pathway. It has been suggested that the two tRNA exons are linked by an unknown tRNA ligase (Weitzer and Martinez 2007). Mice deficient in Clp1 PNK activity develop motor neuron disease, resulting from motoneuron cell, which is attributed to the accumulation of tRNA fragments caused by impaired pre-tRNA splicing (Hanada et al. 2013). A human genetic disease caused by a mutation in the *Clp1* gene has also been reported and presents as a neurological disease (Karaca et al. 2014; Schaffer et al. 2014).

Instead of Clp1, multi-domain enzymes with several catalytic functions are required in the reaction processes of the 5′-phosphate ligation pathway. For example, the fungus *Saccharomyces cerevisiae* protein Trl1 (*Sc*-Trl1) (Apostol et al. 1991), the plant *Arabidopsis thaliana* protein Rnl (*At*-Rnl) (Englert and Beier 2005), and the protist *Trypanosoma brucei* protein Trl1 (*Tb*-Trl1) (Lopes et al. 2016) are multi-domain enzymes with an adenylyltransferase/ligase domain in the N-terminal region, a PNK domain in the central region, and a cyclic phosphodiesterase domain in the C-terminal region. Interestingly, *S. cerevisiae* Clp1 (*Sc*-Clp1) has no PNK activity (Ramirez et al. 2008), and *Sc*-Trl1 functions in the pre-tRNA splicing reaction (Apostol et al. 1991). The amphioxus *Branchiostoma floridae* has two separate enzymes, adenylyltransferase/ligase and PNK/CPDase, which are thought to co-operate in the 5′-phosphate ligation pathway (Englert et al. 2010). In contrast, the tRNA ligase RtcB/HSPC117 enzyme has been identified in the 3′-phosphate ligation pathway, which is widely conserved among bacteria (Tanaka et al. 2011), archaea (Englert et al. 2011), and eukaryotes (Popow et al. 2011). In terms of the other function of Clp1, in mRNA 3′-end formation, the Clp1s of *S. cerevisiae*, *Homo sapiens*, and *A. thaliana* are involved in mRNA 3′-end cleavage and as components of the polyadenylation factor complex (Minvielle-Sebastia et al. 1997; de Vries et al. 2000; Xing et al. 2008). Notably, mRNA 3′-end formation does not require the PNK activity of Clp1 and occurs even when PNK is inactivated, as in the case of *Sc*-Clp1 (Gross and Moore 2001; Ramirez et al. 2008). Therefore, Clp1 is an enzyme that mainly targets tRNAs and mRNAs.

Eukaryotic Clp1 family proteins consist of two major groups with different biological functions: the Clp1 group mentioned above and the Nol9/Grc3 group (Braglia et al. 2010). Here, the enzymes in the latter group are designated Nol9 in Metazoa and Plantae and as Grc3 in Fungi, and both the Nol9 and Grc3 proteins play important roles in precursor ribosomal RNA (pre-rRNA) processing. For example, *S. cerevisiae* Grc3 (*Sc*-Grc3) and *H. sapiens* Nol9 (*Hs*-Nol9) interact with endoribonuclease LAS1 (LAS1L in mammals) and are involved in rRNA biogenesis through the cleavage of the internal transcribed spacer 2 (ITS2) and have PNK activity against 26 rRNA in *S. cerevisiae* (Castle et al. 2013) and against 28S rRNA in *H. sapiens* (Heindl and Martinez 2010; Gordon et al. 2019). The PNK activity *Sc*-Grc3 is also involved in transcription termination by RNA polymerase I (Braglia et al. 2010). Therefore, the Clp1 family proteins are some of the important factors that process tRNA, mRNA, and rRNA, the three typical RNAs that control genetic information.

Clp1 proteins and/or enzymes with Clp1-like PNK domains (Clp1_P) have been previously reported in eukaryotes and archaea (Noble et al. 2007; Jain and Shuman 2009). We have analysed full-length genomes in the three domains of life and reported that the Clp1 family proteins are present in a wide range of phylogenetically diverse but restricted species of bacteria and exert PNK activity against RNA (Saito et al. 2019). Bacterial Clp1 has an essentially simple domain structure, in which the Clp1_P domain is important for its PNK activity. In archaeal Clp1, there is a conserved domain in the C-terminal region. By comparison, eukaryotic Clp1 is essentially composed of three domains, the Clp1_P domain in the central region, and N-terminal and C-terminal domains on either side of it (Noble et al. 2007). Although several conserved domains have been suggested to occur on both sides of the central Clp1_P domain in Nol9 and Grc3 (Pillon et al. 2017; Gordon et al. 2019; Saito et al. 2019), no detailed analysis of the domain structures across a wide range of lineages has been undertaken.

In light of the studies discussed above, in this study, we precisely analysed the molecular evolution of the Clp1 enzymes in eukaryotes as a whole, to clarify how they evolved from prokaryotic Clp1, which has a simple structure, into diverse molecules in eukaryotes and how PNK-inactive forms of Clp1, such as *Sc*-Clp1, have evolved. We performed a large-scale molecular evolutionary analysis of the Clp1 molecular species and their protein domains using 358 full-length genomes of eukaryotic species. In so doing, we identified a group of protist proteins we consider to be the origin of the Clp1 family proteins that arose during eukaryotic evolution. An analysis of PNK activity of recombinant enzymes encoded by the three *Clp1* genes of *T. brucei*, a representative species in the phylum Euglenozoa, suggested that the Clp1 group proteins, which are thought to have arisen by gene duplication, include both PNK-active and PNK-inactive forms, and that *Sc*-Clp1 may have evolved from the PNK-inactive Clp1 lineage. In the fish infraclass Teleostei and the plant phylum Magnoliophyta, which are presumed to have undergone multiple *Clp1* gene duplications associated with whole-genome duplications, the amplified *Clp1* gene has been partially modified and diversified by domain gain or deletion during evolution. In these species and some other vertebrates, including humans, the creation of protein isoforms has occurred via alternative splicing. In this paper, we summarise these findings and propose an integrated molecular evolutionary model of the origin and diversification mechanisms of the Clp1 family proteins in eukaryotes.

## Results and Discussion

### Comprehensive Analysis of Clp1 Family Proteins in Complete Eukaryotic Genomes

In our previous paper, we examined the distribution of the Clp1 family proteins in the complete genomes of 288 eukaryotes (Saito et al. 2019). However, no detailed analysis of the abundance of the Clp1 family proteins in each eukaryotic species or the classification of the Clp1 family protein types was performed. In this study, using known Clp1 sequences and those of its fellow family member proteins Nol9/Grc3 as query sequences (Supplementary Table S1), we performed a comprehensive sequence similarity search (E-value ≤ 1e−4) for each coding sequence on the 358 complete genomes registered in the RefSeq database (as of September 2020). Clp1 family proteins were detected in 1,264 of the 12,333,502 sequences searched, and we confirmed that they are conserved in most species of eukaryotes (350/358; 97.8%) (Supplementary Table S2A), whereas they do not occur the protists Cryptophyta and Rhizaria. Among these 1,264 sequences, Clp1 sequences annotated as splicing isoforms were present in higher organisms such as Metazoa and Viridiplantae, demonstrating that the Clp1 family proteins have diversified through alternative pre-mRNA splicing in these organisms. We then calculated the number of Clp1 family proteins, excluding splicing variants, in each species as the number of independent *Clp1* genes per species at this stage. The results showed that 798 proteins were encoded in the 350 species that had at least one Clp1 family protein (2.3 ± 1.0 proteins per species on average; Supplementary Table S2A). Approximately 10% of Metazoa, approximately 40% of Viridiplantae, and 100% of Euglenozoa contained three or more Clp1 family proteins in each species (Figure 1A). Approximately 90% of Euglenozoa species encoded four Clp1 family proteins. In contrast, in some taxa outside the Euglenozoa, Metazoa, and Amoebozoa, only one Clp1 family protein was present in each species, probably as the result of gene loss (Figure 1A).

**Fig. 1.**
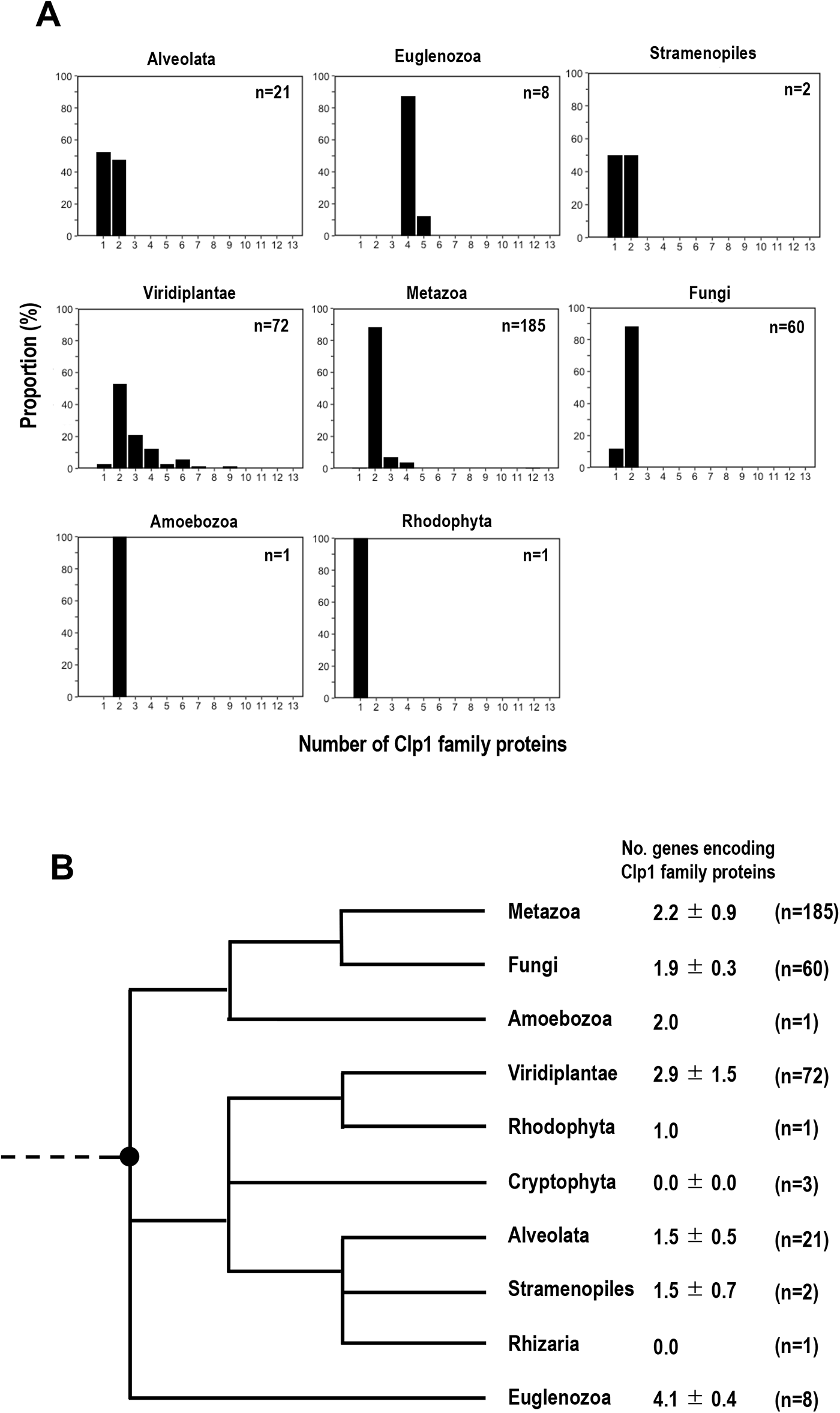
Numbers of Clp1 family proteins differ among eukaryotic taxa. (A) The distribution of the number of Clp1 family proteins in each taxon is shown. Taxon names are shown above each figure (see also Table 1). “n” indicates the number of species used in the analysis. (B) Average numbers (±standard deviations) of *Clp1* genes are mapped against the evolutionary phylogenetic tree of eukaryotes of Adl et al. (2019) (modified). Because the origin of the eukaryotes is unclear on this phylogenetic tree, it is indicated by a dotted line. The black circle indicates the possible time point at which gene duplication is presumed to have occurred.

**Table 1.**
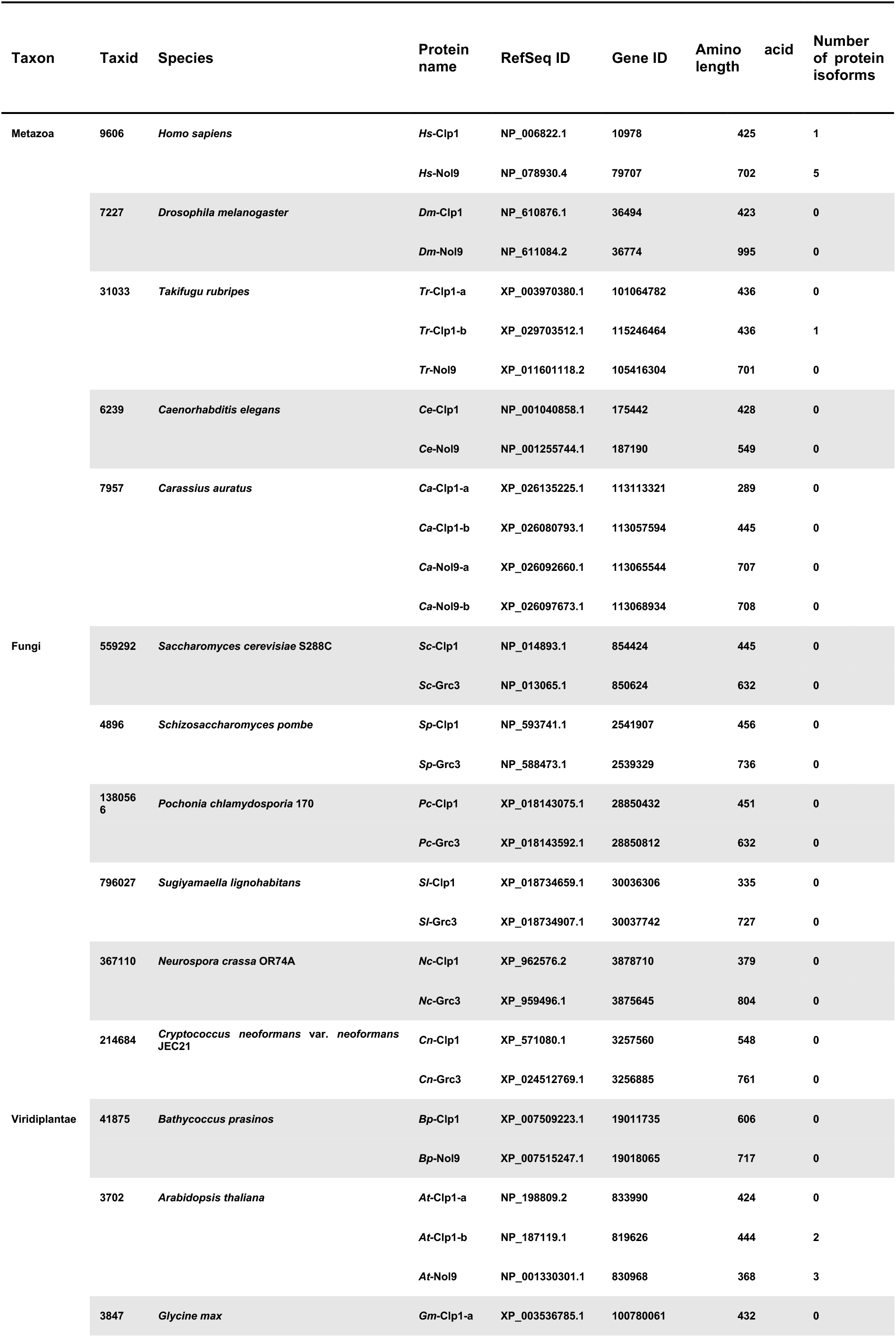
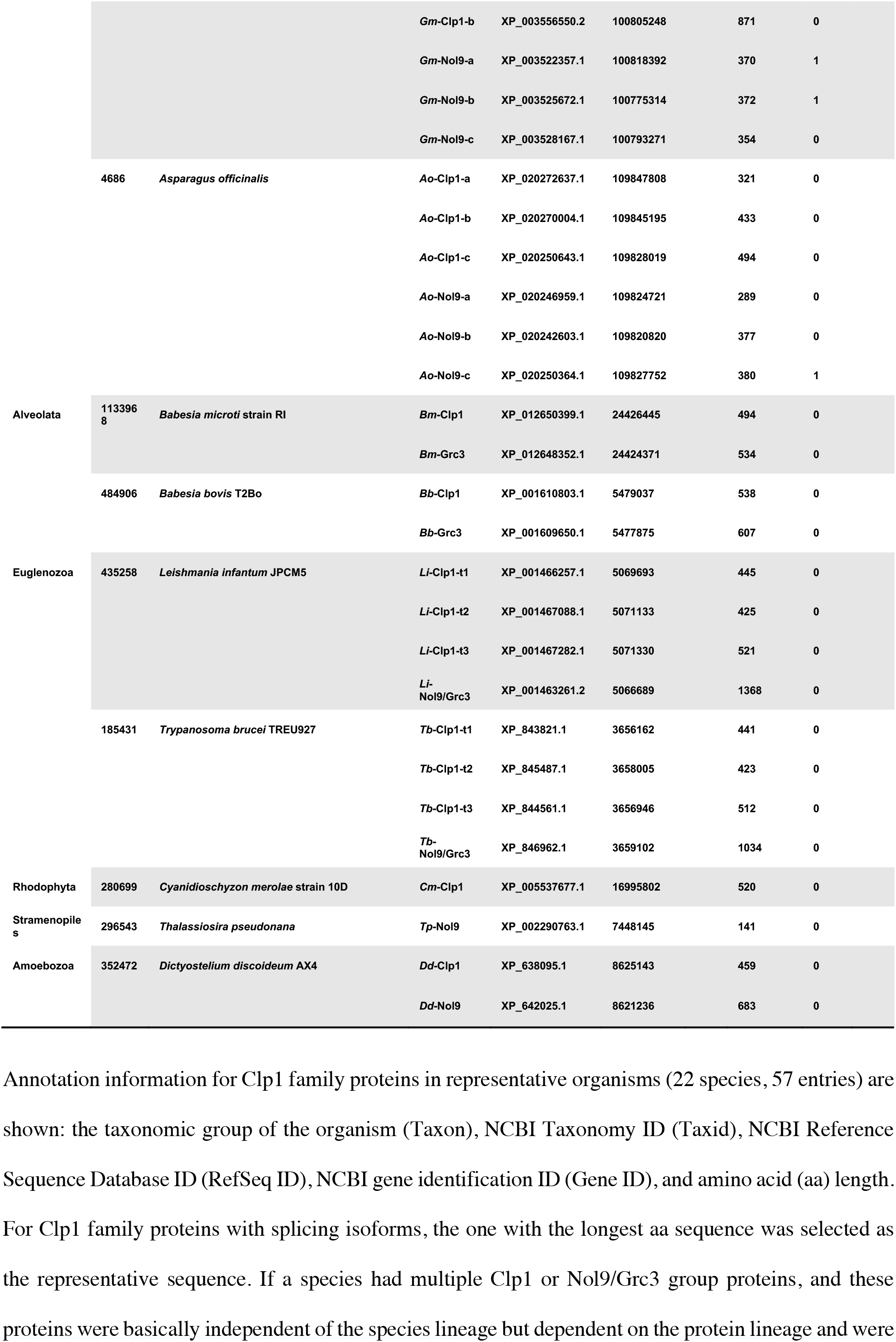
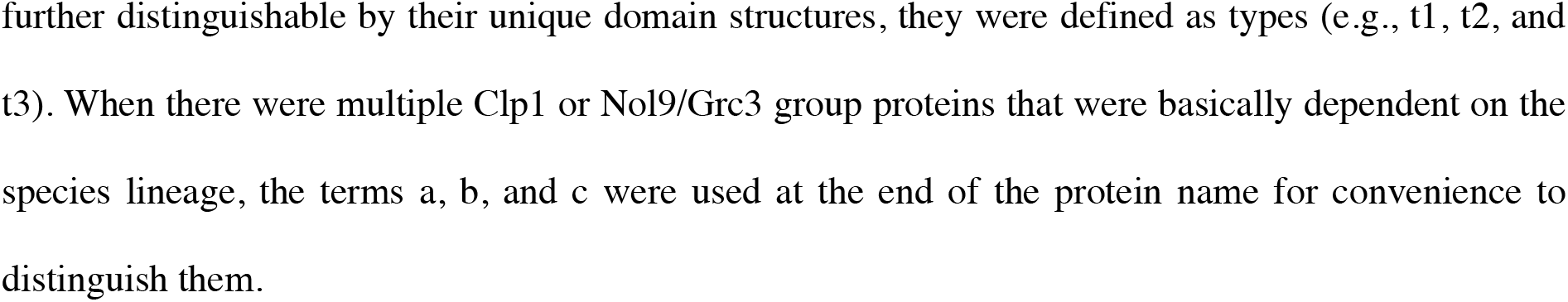
Classification of Clp1 family proteins of representative eukaryotic species.

To classify the Clp1 family proteins present in multiple species in detail according to their sequence similarities, 254 Clp1 family proteins, including the representative sequences in Table 1, were selected from the 1,264 sequences, and a molecular evolutionary phylogenetic tree was constructed from their full-length sequences. The resulting phylogenetic tree was largely divided into two groups: the Clp1 and Nol9/Grc3 group proteins (Supplementary Figure S1). In 22 representative species (Table 1), the classification of the Clp1 family proteins showed that species such as *H. sapiens* (Metazoa), *S. cerevisiae* (Fungi), and *Babesia microti* (Alveolata) had one each of the Clp1 and Nol9/Grc3 group proteins. When a species had two or more Clp1 group proteins or Nol9/Grc3 group proteins, and each protein with unique characteristics domain formed a clade, regardless of the species, it was defined as a “type”. The “types” will be described later in detail, together with the domain structures. In other cases, a, b, or c was added to the end of the protein name for convenience to distinguish multiple proteins. For example, *Asparagus officinalis* (Viridiplantae) encodes three Clp1 group proteins and three Nol9/Grc3 group proteins, so we designated the former *Ao*-Clp1-a, *Ao*-Clp1-b, and *Ao*-Clp1-c, and the latter *Ao*-Nol9-a, *Ao*-Nol9-b, and *Ao*-Nol9-c (Table 1). Among the representative species, *A. officinalis* had the largest total number of Clp1 family proteins (six). In contrast, only one Clp1 was encoded in *Cyanidioschyzon merolae* (Rhodophyta) and one Nol9 in *Thalassiosira pseudonana* (Stramenopiles). The lengths of the protein sequences sometimes differed in each species. For example, the two Clp1 group proteins of *Takifugu rubripes* (Metazoa), *Tr*-Clp1-a and *Tr*-Clp1-b, have exactly the same length (436 amino acid [aa]), whereas the two Clp1 group proteins of *Glycine max* (Viridiplantae), *Gm*-Clp1-a and *Gm*-Clp1-b, are 432 and 871 aa, respectively, an approximately two-fold difference. The regions of the protein that reflect these length differences are discussed below, together with the analysis of the protein domain structures.

Because differences in the numbers of Clp1 family proteins were observed among taxonomic groups, we mapped the number of Clp1 family proteins against the phylogenetic tree of the eukaryotes (Adl et al. 2019) to clarify when the Clp1 family proteins were acquired or lost during evolution. In Figure 1B, the number of Clp1 family proteins is indicated as the mean (±standard deviation) for each taxonomic group. The number of Clp1 family proteins varied from 0 to approximately 4, depending on the eukaryotic taxon, and no clear phylogenetic increase or decrease was detected. It is generally accepted that in eukaryotes, important genes have been genetically duplicated during evolution and that the proteins they encode have also diversified (Marques et al. 2008; Rivera and Swanson 2022). However, variations in the number of Clp1 family proteins among these taxa suggest that the numbers of these proteins have increased or decreased non-phylogenetically. However, we have previously reported that a limited number of prokaryotic species (approximately 1% of bacteria and 40% of archaea) encode Clp1 family proteins (Saito et al. 2019). In that study, most species with a Clp1 family protein had only one protein (average in Bacteria 1.0, and average in Archaea 1.1). In contrast, the present study shows that the average number of Clp1 family protein in eukaryotes is 2.3 ± 1.0, suggesting that the gene encoding this protein duplicated at least once in the eukaryotic root organism (dotted line in Figure 1B).

### Diversification of Clp1 Group Proteins and Their Domain Structures in Eukaryotes

The evolution of the Clp1 family proteins was then analysed at the level of their functional domains. We used two methods to estimate the conserved domain regions in the 254 representative protein sequences (Supplementary Table S2B) used in the phylogenetic analysis described above: a search against the Pfam-A domain database and a sequence similarity search. The results are shown in Figure 2 (domain structures of 110 Clp1 group proteins) and Figure 3 (domain structures of 144 Nol9/Grc3 group proteins) which present the overall Clp1 phylogenetic tree in two parts. In our previous study, we reported that the Clp1 group proteins are composed of three domains, the Clp1_P domain in the middle of the protein, which is essential for PNK activity, and the Clp1_eN and Clp1_eC domains in N-terminal and C-terminal regions, respectively (Saito et al. 2019). Here, we report that the N-terminal domain of *Caenorhabditis elegans* Clp1 (*Ce*-Clp1) is required for ATP binding, and the C-terminal domain of *Ce*-Clp1 may be involved in the stability of the PNK domain (Dikfidan et al. 2014). A search against the domain database showed that, except for the protists, the Clp1 proteins in the taxa Metazoa, Fungi, and Viridiplantae have three domains (Clp1_eN, Clp1_P, and Clp1_eC). However, in the protists (Euglenzoa and Alveolata), which have not been analysed in previous studies, although the Clp1_eN and Clp1_P domains were detected, no domain was detected in the C-terminal region. Therefore, we performed a sequence similarity search (E-value of ≤ 1e−4 and query coverage of ≥ 30%) with BLASTP using protein regions that were not identified in the domain search. We then confirmed the conservation of the hit sequences with an aa alignment and defined them as new domains. For example, the average length of each newly defined domain was 144.3 ± 9.7 aa for Clp1_euC1, 205.7 ± 30.4 aa for Clp1_alC, and 77.2 ± 36.2 aa for Nol9_eN3 (Figures 2 & 3). In this way, we found that different C-terminal domains were conserved in the Euglenzoa and Alveolata Clp1 group proteins: the C-terminal domain was replaced by the Clp1_euC1, Clp1_euC2, or Clp1_euC3 domain in Euglenzoa and by the Clp1_alC domain in Alveolata (Figure 2, see also Supplementary Tables S3–S4 & Supplementary Figures S2–S6). Here, the three Clp1 group proteins with different C-terminal regions in Euglenozoa each formed a clade across species and were defined as types 1–3. An aa sequence alignment of all 25 Euglenzoa Clp1 group proteins showed that there were basically three proteins of each type in each of the eight Euglenozoa species (Supplementary Figure S2 & see also Supplementary Table S3). Among these eight species, *Leishmania mexicana* contained two instances of type 3 (Supplementary Table S3). In summary, 71 of the 107 proteins (66%) used in Figure 2 are derived from Metazoa, Fungi, Viridiplantae and are composed of three domains, as reported previously, indicating that these domains play important roles in a wide range of eukaryotic species. Exceptionally, a small number of proteins were found to lack some of the three domains. For instance, 27 of the 107 proteins (25%) derived from protists (Euglenzoa and Alveolata) had a structure in which only the C-terminal region was replaced by another domain.

**Fig. 2.**
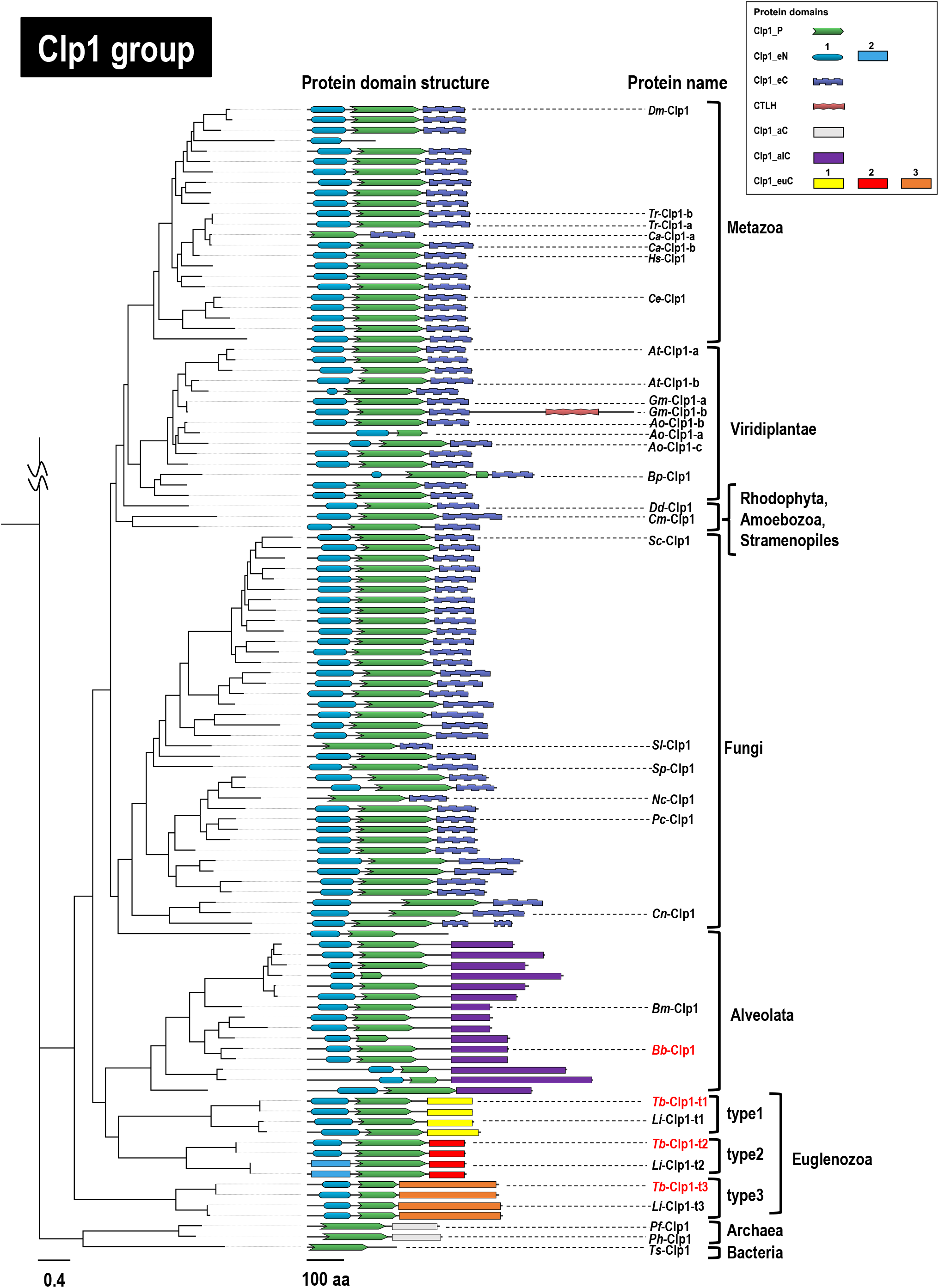
Phylogenetic relationships among Clp1 family proteins (Clp1 group) and their protein domain structures. A phylogenetic tree constructed from a total of 110 Clp1 group protein sequences is shown (Supplementary Table S2B). The 110 Clp1 group sequences consist of 107 protein sequences from 94 eukaryotes, two protein sequences from two archaea, and one protein sequence from a bacterium. Each full-length amino acid sequence of the Clp1 group proteins was used for the phylogenetic analysis, and midpoint rooting was applied during tree visualisation. The entire molecular evolutionary phylogenetic tree of the Clp1 family proteins is shown in Supplementary Figure S1. The scale bar under the tree indicates the number of amino acid substitutions per site. Symbol for protein domain structures are summarised in the box. Protein domains were identified using two methods: (i) searches against the Pfam database; and (ii) manual amino acid sequence alignments (all symbols are rectangles). Protein names of the query sequences used to detect the protein domain structures are indicated in red letters (see Supplementary Figures S3–S6). See also Supplementary Table S4 for details of domain structures.

**Fig. 3.**
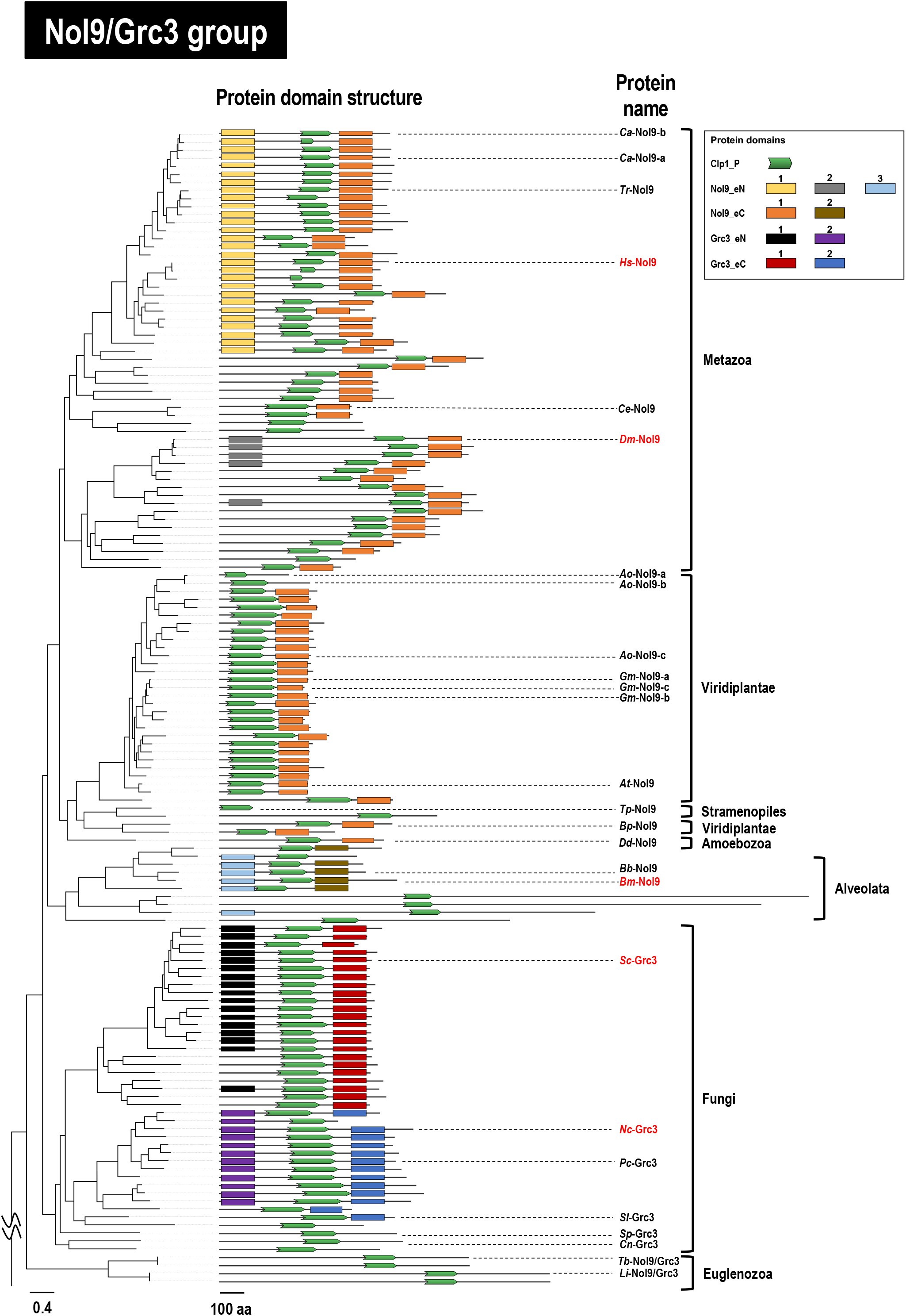
Phylogenetic relationships among Clp1 family proteins (Nol9/Grc3 group) and their protein domain structures. A phylogenetic tree constructed from a total of 144 Nol9/Grc3 group protein sequences from 133 eukaryotes is shown (Supplementary Table S2B). Each full-length amino acid sequence of the Nol9/Grc3 group proteins was used for the phylogenetic analysis. Protein names of the query sequences used to detect protein domain structures are indicated in red letters (see Supplementary Figures S7–S12). See also the legend to Figures 2 for other details.

Next, we wanted to explain the domain structure of *A. officinalis* Clp1 proteins such as *Ao*-Clp1-a, *Ao*-Clp1-b, and *Ao*-Clp1-c shown in Table 1. As shown in Figure 2, the Clp1 group proteins are mainly composed of three domains (Clp1_eN, Clp1_P, and Clp1_eC). Although two proteins of *A. officinalis*, *Ao*-Clp1-b (433 aa) and *Ao*-Clp1-c (494 aa), have the same three-domain structure, *Ao*-Clp1-a (321 aa), which is the shortest of the three proteins, has only two domains, Clp1_eN and Clp1_P, showing that multiple proteins within a species have diversified by changing their domain structures. Similar examples, although not numerous, were also found in in *Carassius auratus* of Metazoa, *Ca*-Clp1-a (289 aa) and *Ca*-Clp1-b (445 aa) (Table 1 & Figure 2), suggesting that multiple proteins are responsible for the limited control required only for the restricted species. In *G. max*, *Gm*-Clp1-a (432 aa) and *Gm*-Clp1-b (871 aa) differ approximately two-fold in their aa sequence lengths. The two proteins share the same three domains, Clp1_eN, Clp1_P, and Clp1_eC, but *Gm*-Clp1-b has a longer total length due to a sequence of approximately 400 aa containing a ‘CTLH domain’ in the C-terminal region. The CTLH domain is found in proteins involved in functions such as microtubule dynamics, cell migration, nucleokinesis, and chromosome segregation and is reported to play a role in protein–protein interactions (Emes and Ponting 2001).

### Nol9/Grc3 Group Proteins are More Diverse in Their Domain Structures than Clp1 Group Proteins

The Nol9/Grc3 group is nominally two groups of proteins, Nol9 found in mammals and plants and Grc3 found in fungi. The *H. sapiens* Nol9 protein (*Hs*-Nol9) is reported to have N- and C-terminal domains in addition to the Clp1_P domain that is essential for PNK activity, as mentioned above (Gordon et al. 2019). In our previous study, we designated these three domains, Nol9_eN, Clp1_P, and Nol9_eC, because they are conserved among the Mammalia (Saito et al. 2019), but because several new domains were detected in the current analysis, we have numbered them systematically and designated them, for instance, Nol9_eN1 and Nol9_eC1 in this paper (Figure 3). In considering the other protein, Grc3, we subjected the fungal *Sc*-Grc3 protein to a functional analysis of its domains. It has been reported that *Sc*-Grc3 is also composed of a Clp1_P domain (Braglia et al. 2010), an N-terminal domain involved in the survival of the yeast, and a C-terminal domain involved in the stability of endonuclease binding (Pillon et al. 2017). In this study, we again attempted to systematically detect novel domains in the N- and C-terminal regions of the Nol9/Grc3 protein group by combining the aforementioned domain database search and a similarity search. We thus identified the Clp1_P domain in all the proteins of the Nol9/Grc3 group and several proteins with unique N-terminal and C-terminal domains in each taxon (Figure 3). However, compared with the Clp1 group, in which the proteins predominantly have three-domain structures (Clp1_eN, Clp1_P, and Clp1_eC), the Nol9/Grc3 group is more diverse, with many proteins lacking either the N- or C-terminal domain (Figure 3 & see also Supplementary Figures S7– S12). Fungal Grc3 proteins are basically composed of three domains, with an N- and C-terminal domain on either side of the Clp1_P domain, and there are two main groups of proteins with this domain structure, depending on the taxonomic group. One Grc3 group consists of three domains, similar to *Sc*-Grc3 of *S. cerevisiae* described above, and 17 (41%) of the 41 fungal species in Figure 3 are classified in Saccharomycotina (Figure 3, see also Supplementary Table S2B & Supplementary Figure S11). The other group, in which the Grc3 proteins have N- and C-terminal domains that differ from those in the previous Grc3 group, accounted for 11 (27 %) of the 41 fungal species, all of which were classified as Pezizomycotina (Figure 3 & see also Supplementary Figure S12). An example of this group is the *Nc*-Grc3 protein of *Neurospora crassa*. In addition to these proteins, other lineage-specific proteins in Fungi consisted solely of the Clp1_P domain or lacked either the N- or C-terminal domain, although the proportions of these proteins were low. We speculated that in these proteins, the N- and C-terminal domains were optimised for each taxon; e.g., they were specific for the substrate specificity of the enzyme or some other function. Figure 3 shows that the lengths of the proteins in the Nol9/Grc3 group varied, so we analysed this variation in detail. First, the mean sequence length (± its standard deviation) of the 144 Nol9/Grc3 group proteins used in Figure 3 was 688 ± 296 aa. The longest was the Nol9 protein (2,440 aa) of *Toxoplasma gondii* (Alveolata), which has extremely long N- and C-terminal regions, in which no domain that is conserved among other species was detected. The shortest was the Nol9 protein (144 aa) of *Thalassiosira pseudonana* (Stramenopiles), which completely lacked the N- and C-terminal domains. In 31 Nol9 proteins of Viridiplantae, the mean length of 409 aa was more than 200 aa shorter than the overall mean value due to a missing N-terminal region. In species in which this domain of Nol9/Grc3 is deleted, the function of Nol9/Grc3 may not necessarily require three domains (Gordon et al. 2019). Interestingly, the Euglenozoa located at the root of the Nol9/Grc3 group was composed exclusively of the Clp1_P domain, with no N- or C-terminal domains detected. Finally, multiple Nol9/Grc3 group proteins occurred within the same species, such as -a, -b, and -c, and the same argument can be made as for the Clp1 group proteins.

### Evolution of the PNK Domain (Clp1_P Domain)

We next analysed how the Clp1 PNK domain (Clp1_P domain) has evolved over time during eukaryotic evolution. We analysed the Clp1_P domain of 253 representative protein sequences, excluding the sequence of *Nasonia vitripennis* of Metazoa (RefSeqID: XP_008207576.1), in which Clp1_eN but no Clp1_P domain was detected. Notably, Clp1 and Nol9/Grc3 proteins with the Clp1_P domain were present in *N. vitripennis*. First, the sequence length of the Clp1_P domain was examined. The mean length (± its standard deviation) of 253 Clp1 family proteins was calculated to be 168 ± 34 aa. The largest was the 220-aa Clp1_P domain of the Clp1 protein (XP_571080.1, 548 aa) of *Cryptococcus neoformans*, which is classified in Fungi, and the smallest was the 55-aa Clp1_P domain of the Nol9 protein (XP_018921794.1, 601 aa) of *Cyprinus carpio*, which is classified in the Metazoa. When we examined the four motifs (Walker A, Walker B, Clasp, and Lid) in the Clp1_P domain of *C. carpio* that are important for PNK activity, we found that the sequences corresponding to Clasp and Rid were completely absent. This was also true of 16 proteins in which the Clp1_P domain was shorter than 130 aa (approximately 6% of the total proteins), suggesting that species with Clp1_P domains significantly smaller than the mean length lack PNK activity.

Surprisingly, a phylogenetic tree constructed from the aa sequences of the Clp1_P domain allowed the Clp1 group proteins and the Nol9/Grc3 group proteins to be distinguished simply by the sequence of the Clp1_P domain alone (Supplementary Figure S13). Because a previous study reported that *Hs*-Clp1 of the Clp1 group has PNK activity during pre-tRNA splicing (Weitzer and Martinez 2007; Ramirez et al. 2008) and *Hs*-Nol9 of the Nol9/Grc3 group has PNK activity during pre-rRNA processing (Heindl and Martinez 2010; Gordon et al. 2019), it is possible that the aa sequence of the Clp1_P domain and its substrate specificity for pre-tRNA or pre-rRNA have co-evolved. Moreover, on the phylogenetic tree based on the full-length sequences of the Clp1 family proteins (Supplementary Figure S1), the taxa in both the Clp1 group and the Nol9/Grc3 group predominantly clustered together, but on the phylogenetic tree based on the Clp1_P domain (Supplementary Figure S13), some of the same taxa were divided into a few phylogenetically distant positions. For example, two Fungi and three Euglenozoa species independently localised to the Clp1 group on the tree. Similarly, in the Nol9/Grc3 group, three Fungi and three Metazoa species also localised independently on the tree. On the phylogenetic tree based on the full-length sequences of the Clp1 family proteins, the order of branches reflected the evolutionary lineage of the species, with Euglenozoa located near the prokaryotes (Supplementary Figure S1). In contrast, the phylogenetic tree based on similarities among the Clp1_P domains only classifies Clp1 type 1 of Euglenozoa near Fungi but classifies Clp1 types 2 and 3 far from Clp1 type 1, near Viridiplantae, which does not reflect the phylogenetic order of evolution (Supplementary Figure S13). Using the full-length regions of *T. brucei* Clp1 types 1–3 from Euglenozoa, we analysed their aa sequence similarities in a round-robin fashion and found only low values of aa identity (24%–34%; Table 2). These results suggest that the Clp1_P domain of each type of Euglenozoa Clp1 has evolved independently and that conserved regions outside the Clp1_P domain are more similar to each other than to conserved regions outside the Clp1_P domain in other Clp1 proteins.

**Table 2.**
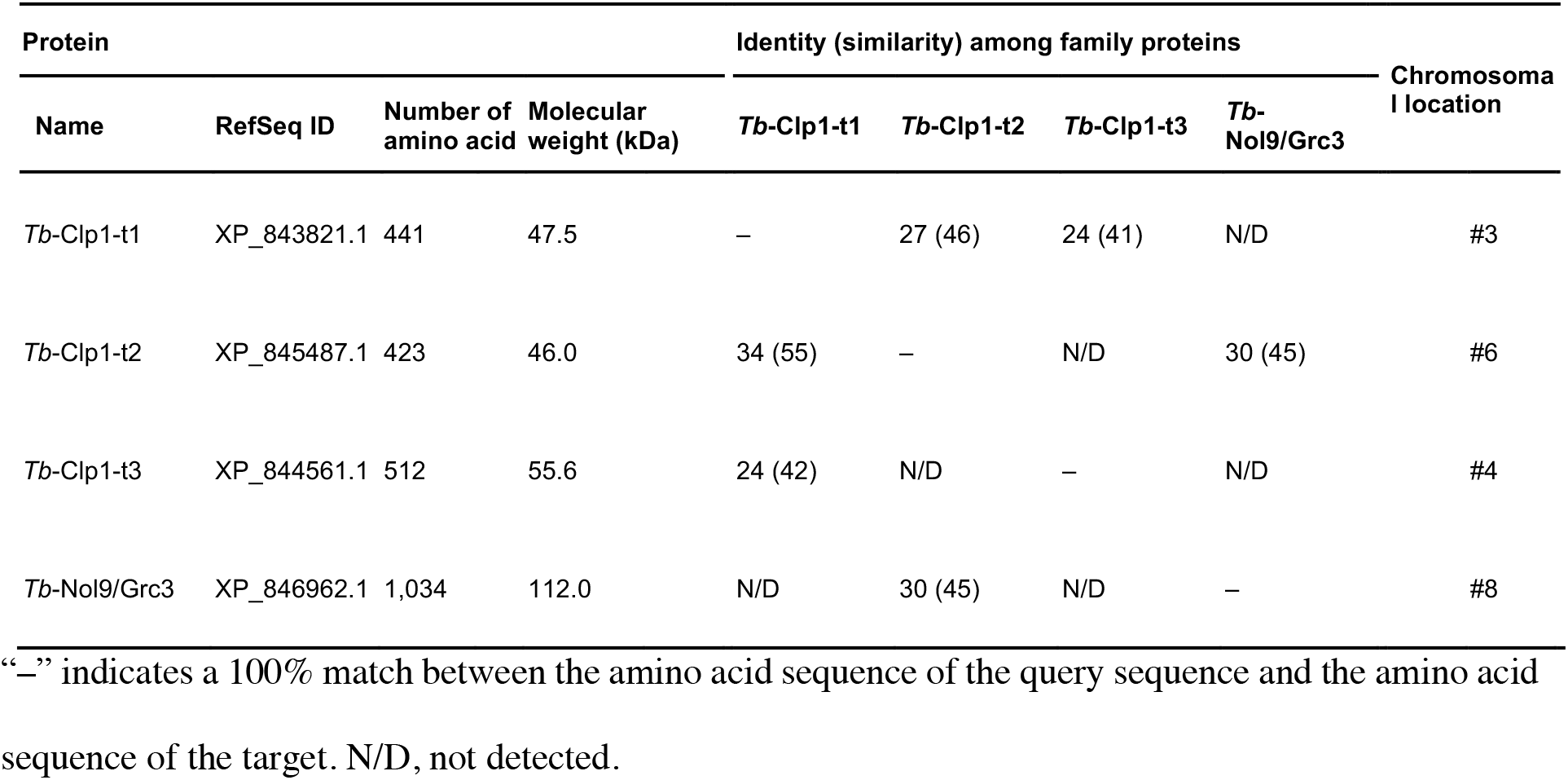
Summary of Clp1 family proteins in T. brucei.

In contrast, on the phylogenetic tree based on the Clp1_P domain, the Clp1 type 2 proteins and Nol9/Grc3 proteins from Euglenozoa localised at the root of each group, suggesting that both proteins are the original forms of each group of proteins in eukaryotes. Moreover, the prokaryotic Clp1 protein localised at the base of the Nol9/Grc3 group, suggesting that prokaryotic Clp1 probably originated as a regulator of pre-rRNA. Therefore, the function of prokaryotic Clp1 in pre-rRNA processing must be investigated in future studies.

### Biochemical Characterisation of the PNK Activities of Three Types of Clp1 Proteins in *T. brucei*

We have described Clp1 types 1–3 in Euglenozoa using a sequence analysis, but their PNK activities must be experimentally verified. To detect and compare the PNK activities of the three Clp1 group proteins of Euglenozoa *T. brucei*, three His-tagged recombinant proteins (*Tb*-Clp1-t1, *Tb*-Clp1-t2, and *Tb*-Clp1-t3) were expressed in *Escherichia coli* and partially purified using TALON® Metal (Cobalt) Affinity Chromatography. The calculated molecular weights of these His-tagged recombinant proteins were 48.6 kDa for *Tb*-Clp1-t1, 47.1 kDa for *Tb*-Clp1-t2, and 56.7 kDa for *Tb*-Clp1-t3, and each size of protein detected by the anti-His-tag antibody was consistent with the size predicted from aa sequence deduced from the corresponding genes (Figure 4 & see also Supplementary Figure S14A– S14C). For example, for *Tb*-Clp1-t2 (47.1 kDa), a partially purified main band was observed slightly below the 50-kDa molecular weight marker (Figure 4A). In a western blotting analysis with an anti-His-tag antibody, this band also has cross-reactivity and peaks in elution fractions #9–#12. These observations were also true for recombinant proteins *Tb*-Clp1-t1 and *Tb*-Clp1-t3 (Figure 4B–4C & see also Supplementary Figure S14B–S14C). These results shows that each of the three recombinant Clp1 group proteins of *T. brucei* was partly purified to a major component on SDS-PAGE using this method. The elution fractions of Clp1 types 1–3 were then examined for PNK activity against single-stranded RNA (ssRNA). PNK activity was only detected in *Tb*-Clp1-t2, and the peak of activity was perfectly consistent with the peak of the *Tb*-Clp1-t2 protein on SDS-PAGE (Figure 4A). In contrast, no clear activity was detected for the remaining two proteins, *Tb*-Clp1-t1 and *Tb*-Clp1-t3 (Figure 4B & 4C). These results clearly indicate that among types 1–3 of the Clp1 group proteins of *T. brucei*, only the recombinant *Tb*-Clp1-t2 protein had PNK activity. A similar experiment was performed with recombinant *Leishmania infantum* protein *Li*-Clp1-t2, which also belongs to type 2, but no PNK activity was detected in this recombinant protein (Supplementary Figure S14D). The consensus sequences of four motifs (Walker A, Walker B, Clasp, and Lid) important for PNK activity (Dikfidan et al. 2014) were examined according to the Clp1 type, and all four motifs were completely conserved in *Tb*-Clp1-t2 (Supplementary Figure S2). All four motifs were also completely conserved in *Tbr*-Clp1-t2 of *T. brucei gambiense*, which is classified in the closely related order Trypanosomatida. In contrast, at least part of the motif sequence in *Li*-Clp1-t2 of *L. infantum* and *Lp*-Clp1-t2 of *L. panamensis*, which are also type 2 Clp1, did not match the consensus sequence (Supplementary Figure S2).

**Fig. 4.**
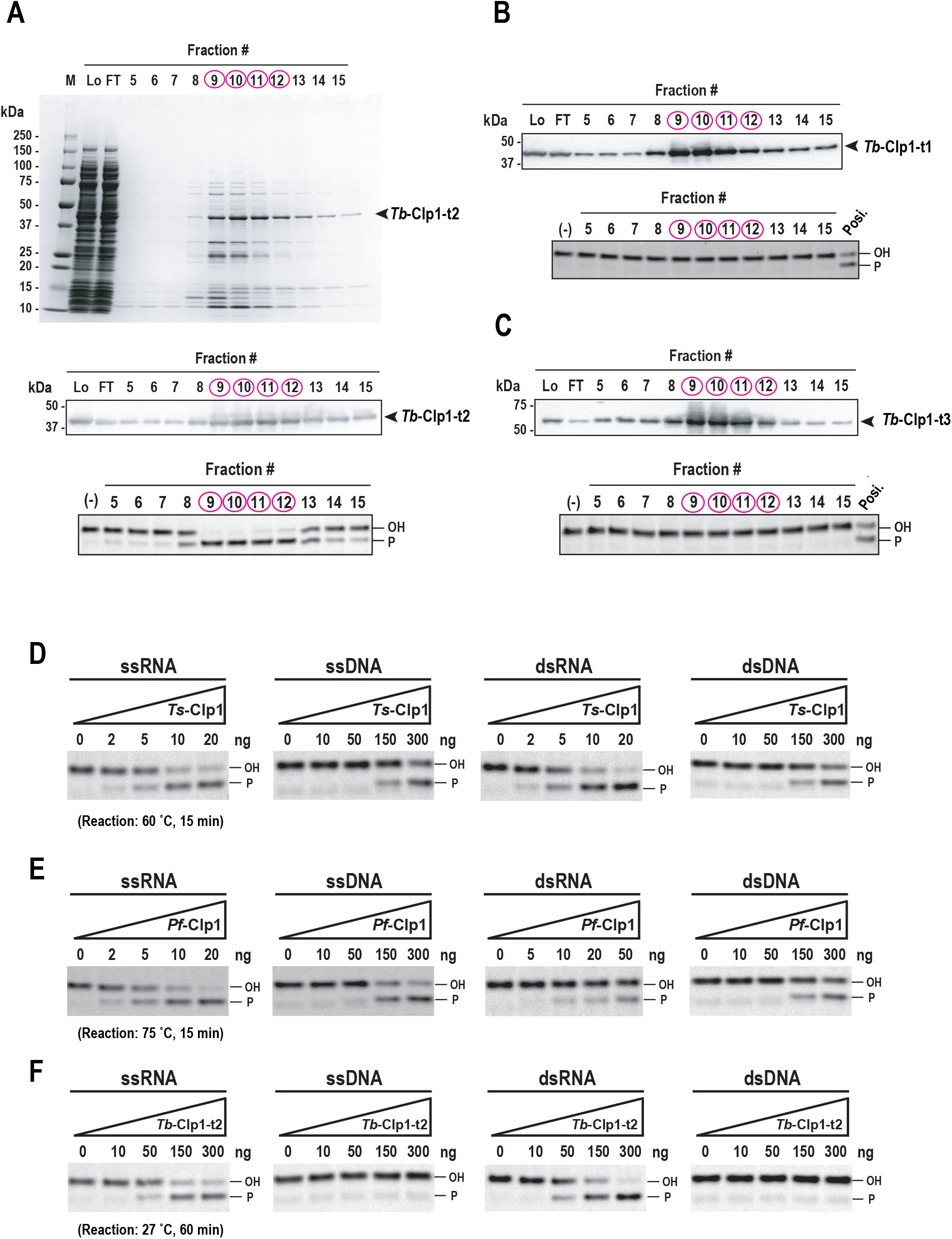
Biochemical characterisation of the recombinant Clp1 group proteins of *T. brucei*. (A) Purification of recombinant *Tb*-Clp1-t2 protein and its PNK activity. 10%–20% SDS-PAGE analysis of the purification of recombinant *Tb*-Clp1-t2 protein using TALON Metal (Cobalt) Affinity Chromatography stained with Coomassie Brilliant Blue (top). Western blotting analysis with an anti-His-tag antibody (middle). PNK activity against single-stranded RNA (ssRNA) as the substrate (bottom). Fractions of the protein peak are indicated by red circles. (B, C) Purification of the recombinant *Tb*-Clp1-t1 and *Tb*-Clp1-t3 proteins (western blotting analysis, top) and their PNK activities (bottom). Arrows indicate the positions of each recombinant protein. Recombinant *Tb*-Clp1-t2 protein was used as the positive control. (D–F) Comparison of PNK activities of three Clp1 proteins: (D) bacterial *Thermus scotoductus* Clp1 (*Ts*-Clp1), (E) archaeal *Pyrococcus furiosus* Clp1 (*Pf*-Clp1), and (F) eukaryotic *Tb*-Clp1-t2 on four substrates (ssRNA, ssDNA, double-stranded RNA [dsRNA], and dsDNA).

We next examined the substrate specificity of *Tb*-Clp1-t2 and compared it with those of prokaryotic archaea *Pyrococcus furiosus* Clp1 (*Pf*-Clp1) and bacteria *Thermus scotoductus* Clp1 (*Ts*-Clp1) (Figure 4D–4F). In a previous study, we reported that *Pf*-Clp1 and *Ts*-Clp1 have a Clp1_P domain, and that the four motifs cited above are highly conserved within this domain. Bacterial *Ts*-Clp1 was used for comparison in this study because the recombinant protein expressed in *E. coli* was found to have PNK activity against DNA and RNA oligonucleotides (Saito et al. 2019). Among the Archaea, *Ph*-Clp1 of *Pyrococcus horikoshii*, in the same genus as *P. furiosus*, has been characterised (Jain and Shuman 2009), and *Pf*-Clp1 and *Ph*-Clp1 are homologous proteins. To compare the reaction conditions at temperatures similar to those *in vivo* for each species, the experiments were conducted at 75 °C for archaeal *Pf*-Clp1, 60 °C for bacterial *Ts*-Clp1, and 27 °C for protist *Tb*-Clp1-t2 (Figure 4F), and reaction time was 15 min for prokaryote Clp1 and 60 min for protist Clp1, so that each change in enzyme activity with the amount of protein used was clear at each temperature. First, prokaryotic *Ts*-Clp1 and *Pf*-Clp1 showed PNK activity against ssRNA and double-stranded RNA (dsRNA) at 5–20 ng per reaction (Figures 4D & 4E). In comparison, PNK activity against ssDNA and dsDNA required about 30 times the amount of enzyme (about 150–300 ng/reaction) of that against ssRNA and dsRNA (Figures 4D & 4E). In a previous study, archaeal *Ph*-Clp1 also showed PNK activity against DNA and RNA and was reported to have stronger activity against RNA than DNA (Jain and Shuman 2009), we confirmed that the experimental results for *Pf*-Clp1 were consistent with the previous report of *Ph*-Clp1. Here, protist *Tb*-Clp1-t2 showed PNK activity against ssRNA and dsRNA at about 50–300 ng/reaction, so it required about 10 times more enzyme and a longer reaction time than prokaryotic *Ts*-Clp1 or *Pf*-Clp1 (Figure 4F). Prokaryotic *Ts*-Clp1 and *Pf*-Clp1 also showed PNK activity against ssDNA and dsDNA when a larger amount of enzyme (approximately 150–300 ng/reaction) was used than against RNA (Figures 4D & 4E), but no activity was detected for *Tb*-Clp1-t2, even when a larger amount was used (300 ng/reaction; Figure 4F). In summary, purified prokaryotic *Ts*-Clp1 and *Pf*-Clp1 showed PNK activity against ssRNA, dsRNA, ssDNA, and dsDNA, although more enzyme was required for the ssDNA and dsDNA substrates. In contrast, purified protist *Tb*-Clp1-t2 was only active against RNA substrates; i.e., ssRNA and dsRNA. These results suggest that the Clp1 group acquired substrate specificity for ssRNA and dsRNA or lost substrate specificity for ssDNA and dsDNA during the early formation of the Clp1 group during the evolution of the protists. Furthermore, compared with prokaryotic Clp1s with their simple domain structures, protist *Tb*-Clp1-t2 and other eukaryotic Clp1s acquired specific domains at their N- and C-termini, respectively (Figure 2). These domains may be involved in the preferential recognition of ssRNA and dsRNA. In previous studies, *Hs*-Clp1 and *Ce*-Clp1 in the Clp1 group showed PNK activity against ssRNA and dsRNA, but not against ssDNA. However, unlike protist *Tb*-Clp1-t2, these enzymes displayed PNK activity against dsDNA (Weitzer and Martinez 2007; Dikfidan et al. 2014). At present, the weaker PNK activities of prokaryotes Clp1s against dsDNA are presumed to have been lost and reacquired in the process of eukaryotic evolution. Therefore, the biological implications of the PNK activity of Clp1 against DNA in nematodes and humans, as well as in prokaryotes, must be verified in future studies.

Finally, we would like to discuss the function of *Tb*-Clp1-t2, the active and early form of eukaryotic Clp1, *in vivo*. The main *in vivo* function of active Clp1 in many organisms is its involvement in pre-tRNA splicing (Ramirez et al. 2008). *T. brucei* has one tyrosine tRNA containing an intron and requires a precise pre-tRNA splicing reaction, catalysed by either an RtcB-dependent pathway that uses the 3′-terminal phosphates of fragmented tRNA exons or a pathway that uses a 5′-terminal phosphate provided by Clp1 of the fragmented tRNA exon (Yoshihisa 2014). In the 5′-phosphate pathway, a tRNA ligase called Trl1 is present in yeast in which the PNK of Clp1 is inactivated (Sawaya et al. 2003; Wang and Shuman 2005). Furthermore, a tRNA ligase called Rnl is present in plants together with a Clp1-similar protein (CLPS3), whose PNK activity has not yet been clarified (Englert and Beier 2005; Xing et al. 2008). Interestingly, a previous study reported that a protein homologous to yeast Trl1 exists in *T. brucei* and functions in joining tRNA exons (Lopes et al. 2016). Therefore, *Tb*-Clp1-t2, whose PNK activity was detected in this study, may function as a backup for Trl1/Rnl or may provide the PNK activity required for RNA repair (Zhang et al. 2012). It should be noted that a previous study suggested that *Tb*-Clp1-t2 is negligibly involved in polyadenylation (Koch et al. 2016).

Remarkably, the protist *T. brucei* is the only representative eukaryote that encodes all of the multiple factors directly involved in joining fragmented tRNA exons, such as RtcB, Trl1/Rnl, and Clp1, although the number of its tRNAs that contain introns is very small (Supplementary Table S5). On the contrary, Metazoa, Fungi, and Viridiplantae, which have more intron-containing tRNAs than protists, lack either RtcB or Trl1/Rnl among the RNA ligase family proteins, although Clp1 is present in each taxon. We presume that an evolutionary event selected the optimal enzyme from Trl1/Rnl and RtcB, depending on the situation of each taxon, which may have included an increased number of intron-containing tRNAs or the inactivation of Clp1.

### Summary of the Molecular Evolution of Clp1 Family Proteins

Based on the large-scale molecular evolution analysis conducted in this study, we propose a model of the diversification of the Clp1 family proteins (Figure 5). When Clp1 originally emerged in prokaryotes, there was basically one Clp1 family protein in each species, with a simple domain structure consisting mainly of the Clp1_P domain and loose substrate specificity. This protein played a role via its PNK activity in the reactions of various RNA and DNA substrates (Figures 2 & 4). In addition to the Clp1_P domain, a C-terminal domain was present in archaeal Clp1 (Saito et al. 2019). In the present study, we found that the Euglenozoa Clp1 family proteins, which are considered to be the most ancient Clp1 proteins in eukaryotes, diversified by gene duplication, giving rise to approximately three Clp1 group proteins (e.g., *Tb*-Clp1-t1, *Tb*-Clp1-t2, and *Tb*-Clp1-t3) and one Nol9/Grc3 group protein (e.g., *Tb*-Nol9/Grc3) in each species. We then speculated on the evolutionary strategies of the three Clp1 group proteins, which are considered to be the starting point of the evolutionary process that culminated in the eukaryotic Clp1 group proteins, based on their PNK activity, substrate specificity, and phylogenetic relationships. We detected no PNK activity in *Tb*-Clp1-t1 and *Tb*-Clp1-t3, at least under the conditions used (Figures 4B & 4C). *Sc*-Clp1 is known to have no PNK activity (Ramirez et al. 2008), and its position on the phylogenetic tree based on the Clp1_P domain shows that it is phylogenetically close to the PNK-inactive *Tb*-Clp1-t1 (Supplementary Figure 13), suggesting that *Sc*-Clp1 was transmitted from the *Tb*-Clp1-t1 lineage. Because *Sc*-Clp1 is involved in pre-mRNA 3′-end formation, which does not require PNK activity (Noble et al. 2007; Ghazy et al. 2012), *Tb*-Clp1-t1 is thought to have a similar function (Clayton and Michaeli 2011). No protein phylogenetically close to *Tb*-Clp1-t3 was found, suggesting that the *Tb*-Clp1-t3 lineage was not acquired by any other organism during the course of evolution or was lost after its acquisition. *Tb*-Clp1-t2, the only PNK active against the RNA substrates, was the earliest member of the eukaryotic Clp1 group on the phylogenetic tree based on the Clp1_P domain (Supplementary Figure 13). As mentioned above, the prokaryotic Clp1 family proteins, which appeared early in the evolution of life are active against both RNA and DNA substrates, whereas *Hs*-Clp1 and *Ce*-Clp1 in Metaozoa have similar substrate specificities to that of *Tb*-Clp1-t2 in that they are mainly active against RNA substrates, suggesting the possible transmission of the *Tb*-Clp1-t2 lineage to this taxon. However, unlike *Tb*-Clp1-t2, both *Hs*-Clp1 and *Ce*-Clp1 have PNK activity against dsDNA, although this point is controversial. Therefore, in the Clp1 group, the aa sequences of the Clp1_P domain and the C-terminal domain of Euglenozoa types 1–3 changed to generate proteins with or without PNK activity or with different functions, thereby increasing the evolutionary options, from which the *Tb*-Clp1-t1 and *Tb*-Clp1-t2 lineages were selected. Therefore, the average number of Euglenozoa Clp1 family proteins is 4.1 ± 0.4, compared with 2.2 ± 0.9 in the more evolved Metazoa and 1.9 ± 0.3 in Fungi, suggesting that the number of Clp1 family proteins in the more evolved taxa may have converged (Figure 1).

**Fig. 5.**
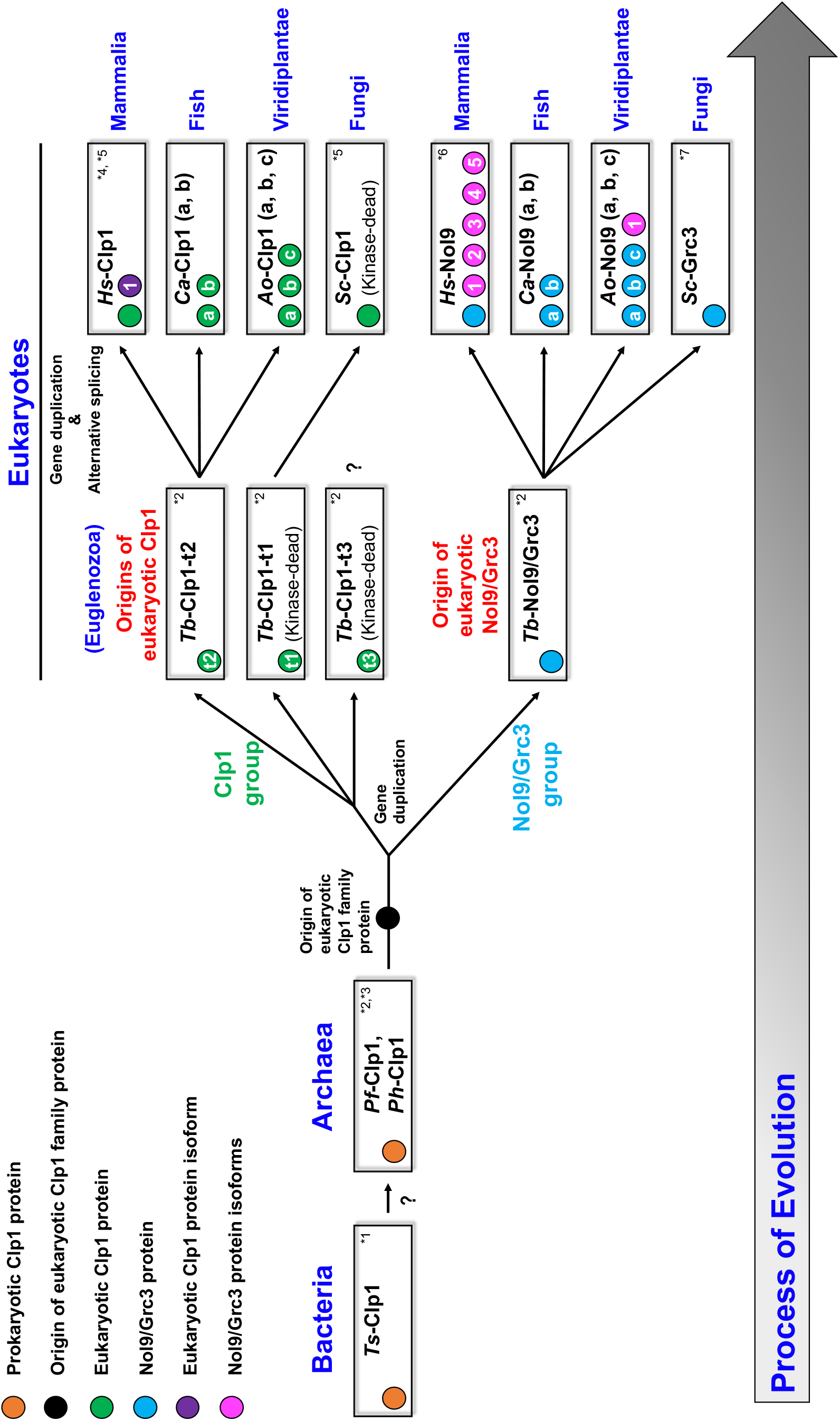
Possible evolutionary model of Clp1 family proteins in the three domains of life. Molecular evolutionary processes of the Clp1 protein family in the three domains of life, based on this and previous studies, are shown. “?” indicates ambiguity. The colour of each circle indicates the protein category. In the circles, “t1–t3” indicate protein types 1–3 of *Tb*-Clp1, “a–c” distinguish individual proteins, and “1– 5” distinguish protein isoforms. References: *1, Saito et al. 2019; *2, this study; *3, Jain and Shuman 2009; *4, Weitzer and Martinez 2007; *5, Ramirez et al. 2008; *6, Heindl and Martinez 2010; *7, Braglia et al. 2010. See text for details.

Having presented a molecular evolutionary model of how the original Clp1 family proteins were selected during eukaryotic evolution, we now explain how the selected proteins diversified. For example, the Clp1 group is basically composed of three domains, Clp1_eN1, Clp1_P, and Clp1_eC, but in Alveolata and Euglenozoa, in addition to the Clp1_eN1, Clp1_eN2, and Clp1_P domains, the C-terminal domain was replaced by the Clp1_alC or Clp1_euC1-euC3 domain in a lineage-specific manner (Figure 2). Similarly, in the Nol9/Grc3 group proteins, the N-terminal and C-terminal domains in some taxa were replaced with unique domains, whereas the Clp1_P domain was usually conserved. Moreover, in Alveolata and Viridiplantae, the Nol9/Grc3 group proteins diversified with changes to their sequence lengths (possibly by including unidentified functional domains; Figure 3). Interestingly, in *H. sapiens*, a vertebrate, the total number of Clp1 family proteins was reduced to two, although two whole-genome duplications are estimated to have occurred (Singh and Isambert 2020). Simultaneously, higher vertebrates produced diverse protein isoforms by alternative splicing (Keren et al. 2010). For example, in *H. sapiens*, there is one protein isoform in the Clp1 group and five in the Nol9/Grc3 group (Figure 5 & Table 1). In the fishes of infraclass Teleostei (Berthelot et al. 2014) and the plant phylum Magnoliophyta (Van de Peer et al. 2017), which both include species with high frequencies of whole-genome duplication, diversity was acquired by the partial modification of the amplified genes and the appearance of protein isoforms, as described above. For example, genes encoding *Ca*-Clp1-a and *Ca*-Clp1-b appeared in the fish *C. auratus*, and genes encoding *Ao*-Nol9-a, *Ao*-Nol9-b, and *Ao*-Nol9-c in the plant *A. officinalis*. These Clp1s are either identical to each other in domain structure or lack some domains (Figures 2 & 3).

## Materials and Methods

### Dataset

To search for Clp1 family proteins in complete genomes, we obtained 358 GenBank files (Supplementary Table S2) annotated as “Complete genome” or “Chromosome” from the Reference Sequence (RefSeq) database (O’Leary et al. 2016) at ftp://ftp.ncbi.nlm.nih.gov/genomes/refseq/; last accessed August 8, 2020. Twenty-two representative species were also selected for a phylogenetic analysis to ensure the inclusion of all taxa of Clp1 family proteins identified in this study (Table 1). For the domain analysis, 18,259 families, together with their reliable annotations, were obtained from the Pfam-A database (version 33.1) (Mistry et al. 2021) at ftp://ftp.ebi.ac.uk/pub/databases; last accessed August 8, 2020.

### Large-scale Identification of Clp1 Family Proteins

To identify Clp1 family proteins in the RefSeq database, a protein–protein BLAST (BLASTP, ver. 2.4.0+) search (Camacho et al. 2009) using the BLOSUM62 matrix (Henikoff and Henikoff 1992) was performed with an E-value of ≤1e−4 and query coverage of ≥30%. As query sequences, we used the aa sequences of eukaryotic Clp1 and its family proteins Nol9 and Grc3, which have already been reported in previous studies (Supplementary Table S1). We used both the full-length and PNK domain regions of each query sequence to comprehensively identify Clp1 family proteins, and each Clp1 family protein set obtained was integrated without duplication (1,264 sequences in total; Supplementary Table S2A). The representative Clp1 family protein set (254 sequences in total; Supplementary Table S2B) was collected as follows. First, 1,264 Clp1 family protein sequences were clustered with 70% sequence similarity using CD-HIT (ver. 4.8.1, March 2019) (Fu et al. 2012) to reduce the number of similar proteins, and sequences were randomly collected from each cluster. We then excluded all protein sequences annotated as splicing isoforms and added the sequences of representative organisms (Table 1) and the sequences of prokaryotes for comparative analysis.

### Amino Acid Sequence Alignment Analysis

The aa sequences of the Clp1 family proteins were aligned using MAFFT L-INS-i ver. 7.394 or 7.471 with the default parameters (Katoh and Standley 2013). The results were visualised using Jalview (ver. 2.11.1.4) (Waterhouse et al. 2009). The colours and scores for alignment conservation are based on the following definitions. Identical aa residues were indicated in blue and partly conserved aa residues in light blue. Gaps (–) were inserted to maximise the number of aa matches. The conservation score for each aa position was indicated as one of 12 ranks (0–11). Partially conserved aa (rank 10) were shown by a plus (+) symbol, and identical aa (rank 11) were indicated by asterisks (*) (Livingstone and Barton 1993). The alignment files created in this step were also used for the phylogenetic tree analysis described in “Molecular phylogenetic analysis” below.

The protein similarity scores among the *T. brucei* Clp1 proteins were calculated using the protein–protein BLAST analysis described in “Large-scale identification of Clp1 family proteins.” Here, “amino acid identity” was defined as the percentage of identical aa residues in two different sequences, and “amino acid similarity” was defined as the ratio of the number of aa that were identical or chemically similar between the query and target sequences (Table 2).

### Molecular Phylogenetic Analysis

For the phylogenetic analysis, we used the above-mentioned alignment file and removed any gaps from it using TrimAL (1.2rev59, 2009) (Capella-Gutierrez et al. 2009). Phylogenetic trees were constructed using the best-fit model, as determined using ModelFinder of IQ-TREE (ver. 2.1.10) (Minh et al. 2020), which is excellent for maximum likelihood analyses of large-scale data. An Ultrafast bootstrap (UFBoot2) analysis (Hoang et al. 2018), which is suitable for processing large numbers of sequences, was performed 1,000 times. The molecular phylogenetic trees were visualised using FigTree (ver. 1.4.4) (http://tree.bio.ed.ac.uk/software/figtree/).

### Domain Analysis of Clp1 Family Proteins

The domain structures of the Clp1 family proteins were estimated using the following two methods. (1) Using the entire aa sequence of each Clp1 family protein as the query, the Pfam-A database (Mistry et al. 2021), in which known domains are registered, was searched using HMMER (ver. 3.2) (Mistry et al. 2013), set to an E-value of ≤1e−3. Their domain structures were visualised using DoMosaics (ver. Rv0.95) (Moore et al. 2014). (2) For protein regions that were not hit in the domain search, a sequence similarity search using BLASTP was performed (E-value of ≤1e−4 and query coverage of ≥30%). The conservation of the obtained sequences was confirmed using an aa alignment (Supplementary Table S3), and they were defined as novel domains (Supplementary Table S4).

### Synthesis of Artificial Genes for Euglenozoan Clp1 Proteins and Construction of Expression Vectors

Four euglenozoan *Clp1* genes (encoding *Tb*-Clp1-t1, *Tb*-Clp1-t2, *Tb*-Clp1-t3, and *Li-*Clp1-t2) (Supplementary Table S6) were used to express recombinant proteins in *E. coli*. First, using a WEB tool (https://www.eurofinsgenomics.jp/jp/service/gsy/orderstart.aspx?type=MyCart) provided by Eurofins Genomics Tokyo, we optimised the nucleotide sequences to match the codon usage of *E. coli*, and synthesised each artificial *Clp1* gene. These synthetic genes were designed to contain *Nde*I and *Xho*I sites at their 5′- and 3′-termini, respectively, and were subcloned into these restriction sites in the pET-23b expression vector (Novagen, Madison, WI, USA). The resulting pET-Clp1 vectors encoded each Clp1 protein with a six-histidine (His) tag at its C-terminal end (Supplementary Table S6).

### Expression and Purification of His-tagged Recombinant Clp1 Proteins

To express the recombinant euglenozoan Clp1 proteins (*Tb*-Clp1-t1, *Tb*-Clp1-t2, *Tb*-Clp1-t3, and *Li*-Clp1-t2), *E. coli* strain BL21(DE3) was transformed with each expression vector containing the artificial euglenozoan *Clp1* genes (Supplementary Table S6). The transformants were pre-cultured in Luria–Bertani (LB) medium containing 50 µg/mL ampicillin for 4 h at 37 °C. Each sample was transferred to 350 mL of LB medium containing the same concentration of ampicillin, incubated at 30 °C for 4 h, and then at 17 °C for 2 h. Isopropyl β-D-1-thiogalactopyranoside (IPTG, 0.4 mM) was added and the cells incubated at 17 °C for a further 14 h to overexpress the desired recombinant Clp1 protein. The cells were harvested by centrifugation (9,000 × g for 3 min at 4 °C), and the protein was extracted using sonication (3–4 min) in His-tag-binding buffer containing 20 mM Tris–HCl (pH 8.0), 500 mM NaCl, 5 mM imidazole, and 0.1% (v/v) NP-40. The insoluble proteins were removed by centrifugation (18,000 × g for 10 min at 4 °C). The recombinant proteins were purified using a TALON Metal (Cobalt) Affinity Resin column (Clontech Laboratories, Inc., Palo Alto, CA, USA) and eluted with a linear gradient of imidazole (0–1,000 mM) in His-tag-binding buffer using the ÄKTA FPLC™ Fast Protein Liquid Chromatography system (GE Healthcare, Princeton, NJ, USA). The eluted protein peak was collected and dialyzed against buffer D containing 50 mM Tris–HCl (pH 8.0), 1 mM ethylenediaminetetraacetic acid (EDTA), 0.02% (v/v) Tween 20, 7 mM 2-mercaptoethanol, and 10% (v/v) glycerol.

The expression and purification of recombinant *Ts*-Clp1 (UniProt AC: E8PQM6) were according to our previous report (Saito et al. 2019). Recombinant *Pf*-Clp1 (UniProt AC: Q8U4H6, 354 aa), a homologue of *Ph*-Clp1 (UniProt AC: O57936, 361 aa), in which PNK activity has been verified (Jain and Shuman 2009), was expressed and purified in the same manner as *Ts*-Clp1.

### Immunoblotting Analysis

For the immunoblotting analysis, purified protein fractions were separated with 10%–20% SDS-PAGE and transferred onto polyvinylidene difluoride (PVDF) membrane (Bio-Rad, Hercules, CA, USA) using a semi-dry blotter (ATTO, Taito-ku, Tokyo, Japan). The blot was incubated with a His-tag-directed monoclonal antibody (MBL, Minato-ku, Tokyo, Japan) dissolved in Can Get Signal Solution 1 (Toyobo, Kita-ku, Osaka, Japan) overnight at 4 °C and washed four times with phosphate-buffered saline (PBS)–0.05% Tween buffer for 5 min each time. It was then incubated with a horseradish peroxidase (HRP)-conjugated goat anti-mouse IgG antibody (Proteintech, Rosemont, IL, USA) dissolved in Can Get Signal Solution 2 (Toyobo) at room temperature for 1 h and washed four times with PBS–0.05% Tween buffer for 7 min each time. Finally, the blot was developed with Western BLoT Chemiluminescence HRP Substrate (Takara, Kusatsu, Shiga, Japan), and the signals were detected using the ChemiDoc XRS+ Imager (Bio-Rad).

### Detection of PNK Activity

PNK activity was assayed by analysing a fluorescein amidite (FAM)-labelled oligoribonucleotide probe on a 15% (w/v) polyacrylamide gel containing 8 M urea, because we have previously demonstrated that the migration of the oligoribonucleotide under these conditions varies with the terminal phosphate structure (Sato et al. 2011; Saito et al. 2019). Basically, the reactions were performed in 20 μL of reaction buffer containing 20 mM Tris–HCl (pH 8.0), 1 mM DTT, 50 mM KCl, 10 mM MgCl_2_, 1 mM ATP, 25 pmol of 5′-R20–FAM-3′, and the purified recombinant euglenozoan Clp1 proteins. After the samples were incubated at 27 °C for 60 min, we added an equal volume of stop solution (8 M urea, 1 M Tris–HCl [pH 8.0], and a small amount of Blue Dextran [Sigma Chemical, St. Louis, MO, USA]) to stop the reactions. The reaction mixtures were heated for 5 min at 70 °C, loaded onto a 15% (w/v) polyacrylamide gel containing 8 M urea, and run for 20 min at 1,600 V and for 90 min at 1,800 V. The reaction products were visualised using the Molecular Imager FX Pro (Bio-Rad). The sequences of the oligoribonucleotides used in this assay are summarised in Supplementary Table S7.

## Acknowledgements

The authors thank Mr. Tomoro Warashina for his technical support during the study and all the members of the RNA Group at the Institute for Advanced Biosciences at Keio University, Japan, for their insightful discussions.

## Funding

This work was supported, in part, by JSPS KAKENHI (grant number JP20J21288, to M.S.), a JSPS KAKENHI Grant-in-Aid for Scientific Research © (grant number JP17K07517, to A.K.), and research funds from the Yamagata Prefectural Government and Tsuruoka City, Japan. The funding bodies played no role in the study design, the data collection or analysis, the decision to publish, or the preparation of the manuscript.

## Data Availability Statements

The data underlying this article are available in the article and in its online supplementary material.

## Notes

### Competing Interest Statement

The authors have declared no competing interest.

