## Supplementary_Information for "Diversification of Polynucleotide Kinase Clp1 Family Proteins and Their Possible Origins in Eukaryotic Evolution"

**Supplementary Table S1. Summary of Clp1 query sequences used in this study**

| Species | Protein name | UniProt AC | Total aa length | Polynucleotide kinase region |
|---|---|---|---|---|
| *Homo sapiens* | Polyribonucleotide 5'-hydroxyl-kinase Clp1 | Q92989 | 425 | 121-307 |
| *Homo sapiens* | Polynucleotide 5'-hydroxyl-kinase Nol9 | Q5SY16 | 702 | 306-470 |
| *Saccharomyces cerevisiae* | mRNA cleavage and polyadenylation factor Clp1 | Q08685 | 445 | 130-335 |
| *Saccharomyces cerevisiae* | Polynucleotide 5'-hydroxyl-kinase Grc3 | Q07845 | 632 | 246-450 |

### Supplementary Table S2. Distribution of Clp1 family proteins in complete eukaryotic genomes

#### (A) Summary of the number of all Clp1 family proteins detected in this study (1,264 proteins).

| | Taxon | Number of CDSs in RefSeq database | Number of Clp1 family protein | Number of Clp1 family protein (except splicing isoform) | Number of species registered | Number of species with Clp1 family protein | Ratio (%) |
|---|---|---|---|---|---|---|---|
| Eukarya | Alveolata | 127,693 | 31 | 31 | 25 | 21 | 84.0 |
| | Amoebozoa | 13,315 | 2 | 2 | 1 | 1 | 100.0 |
| | Cryptophyta | 1,695 | 0 | 0 | 3 | 0 | 0.0 |
| | Euglenozoa | 67,115 | 33 | 33 | 8 | 8 | 100.0 |
| | Fungi | 477,076 | 114 | 113 | 60 | 60 | 100.0 |
| | Metazoa | 8,320,001 | 749 | 406 | 185 | 185 | 100.0 |
| | Rhizaria | 344 | 0 | 0 | 1 | 0 | 0.0 |
| | Rhodophyta | 4,803 | 1 | 1 | 1 | 1 | 100.0 |
| | Stramenopiles | 22,081 | 3 | 3 | 2 | 2 | 100.0 |
| | Viridiplantae | 3,299,379 | 331 | 209 | 72 | 72 | 100.0 |
| | Total | 12,333,502 | 1,264 | 798 | 358 | 350 | 97.8 |

#### (B) Summary of representative Clp1 family proteins used in the phylogenetic analysis (*).

| | Taxon | Number of Clp1 group protein | Number of species with Clp1 group protein | Number of Nol9/Grc3 group protein | Number of species with Nol9/Grc3 group protein | Number of Clp1 family protein | Number of species with Clp1 family protein |
|---|---|---|---|---|---|---|---|
| Eukarya | Alveolata | 15 | 15 | 10 | 10 | 25 | 16 |
| | Amoebozoa | 1 | 1 | 1 | 1 | 2 | 1 |
| | Cryptophyta | 0 | 0 | 0 | 0 | 0 | 0 |
| | Euglenozoa | 12 | 6 | 4 | 4 | 16 | 6 |
| | Fungi | 39 | 39 | 41 | 41 | 80 | 46 |
| | Metazoa | 23 | 20 | 55 | 53 | 78 | 57 |
| | Rhizaria | 0 | 0 | 0 | 0 | 0 | 0 |
| | Rhodophyta | 1 | 1 | 0 | 0 | 1 | 1 |
| | Stramenopiles | 1 | 1 | 2 | 2 | 3 | 2 |
| | Viridiplantae | 15 | 11 | 31 | 22 | 46 | 25 |
| Archaea | Euryarchaeota | 2 | 2 | 0 | 0 | 2 | 2 |
| Bacteria | Deinococcus-Thermus | 1 | 1 | 0 | 0 | 1 | 1 |
| | Total | 110 | 97 | 144 | 133 | 254 | 157 |

(*) To create a sequence set of 254 representative Clp1 proteins, similar sequences were removed from the 1,264 detected Clp1 family proteins with CD-HIT. We also excluded the protein sequences of splicing variants. We then added the sequences of representative organisms (Table 1) and the sequences of prokaryotes for comparative analysis.

**Supplementary Table S3. List of Clp1 family proteins (both Clp1 and Nol9/Grc3 groups) used for amino acid sequence alignments**

| Taxon | Taxid | Species | Protein name | RefSeq ID | aa length |
|---|---|---|---|---|---|
| Euglenozoa | 185431 | *Trypanosoma brucei* TREU927 | *Tb*-Clp1-t1 | XP_843821.1 | 441 |
| | | | *Tb*-Clp1-t2 | XP_845487.1 | 423 |
| | | | *Tb*-Clp1-t3 | XP_844561.1 | 512 |
| | 679716 | *Trypanosoma brucei gambiense* DAL972 | *Tbr*-Clp1-t1 | XP_011772180.1 | 441 |
| | | | *Tbr*-Clp1-t2 | XP_011774153.1 | 423 |
| | | | *Tbr*-Clp1-t3 | XP_011773013.1 | 512 |
| | 420245 | *Leishmania braziliensis* MHOM/BR/75/M2904 | *Lb*-Clp1-t1 | XP_001565679.1 | 445 |
| | | | *Lb*-Clp1-t2 | XP_001566862.1 | 425 |
| | | | *Lb*-Clp1-t3 | XP_001567033.1 | 522 |
| | 5679 | *Leishmania panamensis* | *Lp*-Clp1-t1 | XP_010699796.1 | 463 |
| | | | *Lp*-Clp1-t2 | XP_010701260.1 | 425 |
| | | | *Lp*-Clp1-t3 | XP_010701413.1 | 522 |
| | 929439 | *Leishmania mexicana* MHOM/GT/2001/U1103 | *Lm*-Clp1-t1 | XP_003876277.1 | 445 |
| | | | *Lm*-Clp1-t2 | XP_003877385.1 | 425 |
| | | | *Lm*-Clp1-t3-1 | XP_003877542.1 | 522 |
| | | | *Lm*-Clp1-t3-2 | XP_003877540.1 | 522 |
| | 347515 | *Leishmania major* strain Friedlin | *Lma*-Clp1-t1 | XP_001683978.1 | 445 |
| | | | *Lma*-Clp1-t2 | XP_001684846.1 | 425 |
| | | | *Lma*-Clp1-t3 | XP_001685005.1 | 522 |
| | 435258 | *Leishmania infantum* JPCM5 | *Li*-Clp1-t1 | XP_001466257.1 | 445 |
| | | | *Li*-Clp1-t2 | XP_001467088.1 | 425 |
| | | | *Li*-Clp1-t3 | XP_001467282.1 | 521 |
| | 5661 | *Leishmania donovani* | *Ld*-Clp1-t1 | XP_003861557.1 | 445 |
| | | | *Ld*-Clp1-t2 | XP_003862954.1 | 425 |
| | | | *Ld*-Clp1-t3 | XP_003863112.1 | 521 |
| Alveolata | 31271 | *Plasmodium chabaudi chabaud* | *Pc*-Clp1 | XP_744232.1 | 590 |
| | 5858 | *Plasmodium malariae* | *Pm*-Clp1 | XP_028860391.1 | 633 |
| | 1120755 | *Plasmodium cynomolgi* strain B | *Pcy*-Clp1 | XP_004221125.1 | 591 |
| | 5855 | *Plasmodium vivax* | *Pv*-Clp1 | XP_001614946.1 | 562 |
| | 1133968 | *Babesia microti* strain RI | *Bm*-Clp1 | XP_012650399.1 | 494 |
| | | | *Bm*-Nol9 | XP_012648352.1 | 534 |
| | 5874 | *Theileria annulate* | *Ta*-Clp1 | XP_953879.1 | 493 |
| | 484906 | *Babesia bovis* T2Bo | *Bb*-Clp1 | XP_001610803.1 | 538 |
| | | | *Bb*-Nol9 | XP_001609650.1 | 607 |
| | 353152 | *Cryptosporidium parvum* Iowa II | *Cp*-Clp1 | XP_626224.1 | 601 |
| | 5866 | *Babesia bigemina* | *Bbi*-Nol9 | XP_012767159.1 | 737 |
| | 1537102 | *Theileria equi* strain WA | *Te*-Nol9 | XP_004831100.1 | 597 |
| Metazoa | 7739 | *Branchiostoma floridae* | *Bf*-Nol9 | XP_035677701.1 | 694 |
| | 8364 | *Xenopus tropicalis* | *Xt*-Nol9 | XP_002933893.2 | 639 |
| | 31033 | *Takifugu rubripes* | *Tr*-Nol9 | XP_011601118.2 | 701 |
| | 7918 | *Lepisosteus oculatus* | *Lo*-Nol9 | XP_015192803.1 | 562 |
| | 93934 | *Coturnix japonica* | *Cj*-Nol9 | XP_015738036.1 | 651 |
| | 7957 | *Carassius auratus* | *Ca*-Nol9 | XP_026092660.1 | 707 |
| | 219594 | *Aythya fuligula* | *Af*-Nol9 | XP_032057582.1 | 634 |
| | 9606 | *Homo sapiens* | *Hs*-Nol9 | NP_078930.4 | 702 |
| | 7757 | *Petromyzon marinus* | *Pm*-Nol9 | XP_032827451.1 | 782 |
| | 7245 | *Drosophila yakuba* | *Dy*-Nol9 | XP_002092281.1 | 1,032 |
| | 7227 | *Drosophila melanogaster* | *Dm*-Nol9 | NP_611084.2 | 995 |
| | 7240 | *Drosophila simulans* | *Ds*-Nol9 | XP_016027901.1 | 1,054 |
| | 7241 | *Drosophila subobscura* | *Dsu*-Nol9 | XP_034652937.1 | 873 |
| | 7159 | *Aedes aegypti* | *Aa*-Nol9 | XP_021699171.1 | 1,034 |
| | 2817044 | *Belonocnema kinseyi* | *Bk*-Nol9 | XP_033215408.1 | 754 |
| Amoebozoa | 352472 | *Dictyostelium discoideum* AX4 | *Dd*-Nol9 | XP_642025.1 | 683 |
| Viridiplantae | 51953 | *Elaeis guineensis* | *Eg*-Nol9 | XP_029119051.1 | 407 |
| | 4615 | *Ananas comosus* | *Ac*-Nol9 | XP_020108251.1 | 394 |
| | 3847 | *Glycine max* | *Gm*-Nol9 | XP_003525672.1 | 372 |
| | 22663 | *Punica granatum* | *Pg*-Nol9 | XP_031406670.1 | 375 |
| Fungi | 4956 | *Zygosaccharomyces rouxii* | *Zr*-Grc3 | XP_002498342.1 | 624 |
| | 322104 | *Scheffersomyces stipitis* CBS 6054 | *Ss*-Grc3 | XP_001386260.2 | 663 |
| | 931890 | *Eremothecium cymbalariae* DBVPG#7215 | *Ec*-Grc3 | XP_003647416.1 | 633 |
| | 4950 | *Torulaspora delbrueckii* | *Td*-Grc3 | XP_003682747.1 | 623 |
| | 1071383 | *Kazachstania naganishii* CBS 8797 | *Kn*-Grc3 | XP_022464617.1 | 655 |
| | 28985 | *Kluyveromyces lactis* | *Kl*-Grc3 | XP_453983.1 | 631 |
| | 559292 | *Saccharomyces cerevisiae* S288C | *Sc*-Grc3 | NP_013065.1 | 632 |
| | 56646 | *Fusarium venenatum* | *Fv*-Grc3 | XP_025587698.1 | 720 |
| | 332648 | *Botrytis cinerea* B05.10 | *Bc*-Grc3 | XP_024547363.1 | 776 |
| | 510516 | *Aspergillus oryzae* RIB40 | *Ao*-Grc3 | XP_023093528.1 | 816 |
| | 759273 | *Colletotrichum higginsianum* IMI 349063 | *Ch*-Grc3 | XP_018164591.1 | 755 |
| | 148305 | *Pyricularia grisea* | *Pg*-Grc3 | XP_030986943.1 | 727 |
| | 367110 | *Neurospora crassa* OR74A | *Nc*-Grc3 | XP_959496.1 | 804 |

**Supplementary Table S4. Summary of detected domains in Clp1 family proteins**

| Domain symbol | Protein domain name | Protein domain details | Pfam ID |
|---|---|---|---|
| | Clp1_P | Clp1 polynucleotide kinase domain in eukaryotes | PF16575 |
| | Clp1_eN1 | Clp1 N-terminal domain in eukaryotes | PF16573 |
| | Clp1_eN2 | Clp1 N-terminal domain in eukaryotes | PF16573 |
| | Clp1_eC | Clp1 C-terminal domain in eukaryotes | PF06807 |
| | CTLH | CTLH/CRA C-terminal to LisH motif domain | PF10607 |
| | Clp1_aC | Clp1 C-terminal domain in archaea | *1 |
| | Clp1_alC | Clp1 C-terminal domain in alveolata | *2 |
| | Clp1_euC1 | Clp1 C-terminal domain in euglenozoan type 1 | *2 |
| | Clp1_euC2 | Clp1 C-terminal domain in euglenozoan type 2 | *2 |
| | Clp1_euC3 | Clp1 C-terminal domain in euglenozoan type 3 | *2 |
| | Nol9_eN1 | Nol9 N-terminal domain in metazoan (1) | *1 |
| | Nol9_eN2 | Nol9 N-terminal domain in metazoan (2) | *2 |
| | Nol9_eN3 | Nol9 N-terminal domain in alveolata | *2 |
| | Nol9_eC1 | Nol9 C-terminal domain in metazoan, viridiplantae and amoebozoa | *1 |
| | Nol9_eC2 | Nol9 C-terminal domain in alveolata | *2 |
| | Grc3_eN1 | Grc3 N-terminal domain in fungi (1) | *2 |
| | Grc3_eN2 | Grc3 N-terminal domain in fungi (2) | *2 |
| | Grc3_eC1 | Grc3 C-terminal domain in fungi (1) | *2 |
| | Grc3_eC2 | Grc3 C-terminal domain in fungi (2) | *2 |

Protein domains were identified using two methods: (i) searches against the Pfam database; and (ii) manual amino acid sequence alignments (all symbols are rectangles). *1, Saito et al. 2019, *2, this study.

**Supplementary Table S5. Summary of Clp1 and RNA ligase family enzymes (Trl1/Rnl and RtcB) in the species examined in this study**

| | | | | tRNA splicing pathway | | |
|---|---|---|---|---|---|---|
| | | | Number of tRNAs with introns | 5'-phosphoate ligation pathway | | 3'-phosphoate ligation pathway |
| Domain | Taxon | Species | | Clp1 | Trl1/Rnl | RtcB |
| Eukarya | Metazoa | *Homo sapiens* | 28 | ○ | - | ○ |
| | Fungi | *Saccharomyces cerevisiae* | 62 | ○ | ○ | - |
| | Viridiplantae | *Arabidopsis thaliana* | 72 | ○ | ○ | - |
| | Euglenozoa | *Trypanosoma brucei* TREU927 | 1 | ○ | ○ | ○ |
| Archaea | Euryarchaeota | *Pyrococcus furiosus* DSM 3638 | 2 | ○ | - | ○ |
| Bacteria | Deinococcus-Thermus | *Thermus scotoductus* SA-01 | 0 | ○ | - | ○ |

Distributions of the Clp1 and RNA ligase family enzymes involved in pre-tRNA splicing in each representative organism are shown. Presence or absence of the corresponding gene is indicated by "○" and "–", respectively. **Method**: To determine the presence or absence of the corresponding genes, we performed a sequence similarity search against the RefSeq database using known gene sequences, with an E-value of ≤1e−4 and query coverage of ≥30%. The query sequences (UniProt AC) were *Escherichia coli* RtcB (P46850), *Homo sapiens* RtcB (Q9Y3I0), *Saccharomyces cerevisiae* Trl1 (P09880), *Trypanosoma brucei* Trl1 (Q38AF2), and *Arabidopsis thaliana* Rnl (Q0WL81). For the eukaryotic RefSeq database, see the Dataset section in the Materials and Methods, and for the prokaryotic RefSeq database, we used the August 2018 dataset (ftp://ftp.ncbi.nlm.nih.gov/genomes/refseq/; last accessed September 17, 2019). The number of intron-containing tRNAs in each species is according to the Genomic tRNA Database (GtRNAdb) (http://gtrnadb.ucsc.edu) (Chan and Lowe 2009).

#### Supplementary Table S6. Nucleotide sequences of the euglenozoan *Clp1* genes optimised for *E. coli* codon usage and the amino acid sequences encoded

*Tb*-Clp1-t1-His Tag (Optimized nucleotide sequence)

ATGAGTGCAATCGAACAAGCGAACTTCCGCCTGCAAGCAGGCAGCGAACTGGTCGTGGTGGTGCCCTATTCAACCGAACGCTGCTCTGCCACACTGAAAGTTGTGAGTGGGGCCGGAGGCCTTG
AACCACGTGTTGATGTGCTGGGTGCGCCGGTCATTGTGGATGTCAGCTACAACCTCCCGCCGGGCATTACCTTCACCGTCTTTACCTGGAGCAACGCCAACATCCGCATTGAAGGGTCAAAACAG
CTGGTACAGAATTGCTTCCGGAGTACTACCCATCCGTTTGCGCGGTCGATTGTTGAGTACCATTGCCTGATTCACAATGCACGCCTTTTAGCGGATAAGCAGGGCCTGTTCGGTCCGATGGTGCTC
ATTTGCGGTCAGAATGATACGGAGAAGCATGCCATTTCGCGTACTCTGAGCTCCTATGCTGCGCGTACCGGCTGGGCTCCGCAGTTTGTGGATCTGGATTGTGGCATGGGTCAGCTGCTGTCCACG
CCAGGTACCGTAGCGGCCTGTGTTTCCGAATGTCCGATGACGTTAGACGAAGATACGTCGACGGGTCCGCTGTCAGTTGCGTTCTTTGTTGGTAGCACAGAACCACAGGTTAAAGGCGTTTCTGGG
GAATGGAACATGTTTGCGCCGTATGTGCACTATTGTCGCTTGCTGTTGAGTTGCGTTTCAGAACGTATTGCACGCCATAAAGGTGGCGCTGGAGGCAGCTCTGGCGCGATTATCGTGTTACCCGAGT
TGCGTGGGAGCAATGGCCTGCTCTTTGTAGTGGACATCATTCGCCAGTTCAACATCAGCCACATCTTATGCGTAGGGGACGACTTCCTGTTCTGTGGCCTGCATGAACGCATTCCACGCTTGCGCG
AACACATGGCTTCCCGTGTAGGTGGAGACATTCGTTTGGATAAACTCTCTGGCTCGCCTCACTTTCCGTCCCCTGATACTCGCACAGAACGCTTAAGCTCGATTATGGAGCGGTACTTCTTTGGTGG
CGGGGCGGTCGACTTGCAACCTAGCGAGATCAACCGCCGCTATGCGAGCATCGAGATCCTGCTGCTGAAAGAAGCCAATGGCCAAGCTGTTGTGAGTCCCGTCGAGCAAGATGCACTCGAAGGA
GTCGTGGGTTGCGTTGGCTCGTTATTTGAATCTAGTGCTATCCATGAGAAAGGCGCCCTTTCACTGGCACCGTTTGCACTTGCGCGTGTGCAGGGTATCGATGCCAATGGTGTCTCCCTGCTGGTG
AGCACCCATTCTGCCATTCCGGAACGTCTGACGATGATTGTAGGTGCGTTTCGTTGGGTCACCTCGCTCGAGCATCACCATCACCATCAC

*Tb*-Clp1-t1-Hisx6 (Amino acid sequence)

MSAIEQANFRLQAGSELVVVVPYSTERCSATLKVVSGAGGLEPRVDVLGAPVIVDVSYNLPPGITFTVFTWSNANIRIEGSKQLVQNCFRSTTHPFARSIVEYHCLIHNARLLADKQGLFGPMVLICGQNDT
EKHAISRTLSSYAARTGWAPQFVDLDCGMGQLLSTPGTVAACVSECPMTLDEDTSTGPLSVAFFVGSTEPQVKGVSGEWNMFAPYVHYCRLLLSCVSERIARHKGGAGGSSGAIIVLPELRGSNGLLFVV
DIIRQFNISHILCVGDDFLFCGLHERIPRLREHMASRVGGDIRLDKLSGSPHFPSPDTRTERLSSIMERYFFGGGAVDLQPSEINRRYASIEILLLKEANGQAVVSPVEQDALEGVVGCVGSLFESSAIHEKG
ALSLAPFALARVQGIDANGVSLLVSTHSAIPERLTMIVGAFRWVTSLELEHHHHHH

*Tb*-Clp1-t2-His Tag (Optimized nucleotide sequence)

ATGAGTTCAAACTGCCGCACGGAAGAGTTCAATTTGGAGCGCAAAGAACTGAAAATTCGCTTCCGCTCTTCCGGCTATGTCGTACTCACCGATGGGGCGGCGACGATCTTTGGTGCTCCACTGAAG
AAGAACACTCGTTACGACTTTTCAATGTGCTCAATCCCAGTGGTTTCTCCGGTTGCGTGTCGGTTGCATGTTGGCGGAGACTTCACTTCGGTGGTGACGTACATTCGCACAGACGTGTATGATATCC
ATGCAGTCCTGGACTTCGCTCGCCATGAGGCGAGCAAACGTGGTGTGCCCAGCCTGAACGACACGAATCCTGGCGGAACAAGCAGCAATAGCCTGAAAGAAGGCCCGTGGGGTCCGCGTGTGC
TGGTAGTTGGCGATGTTAATACGGGCAAAAGCTCCCTGTGCCGCAGTCTGGCCAACATGGCGGTGGCCTCGCAAGTGCACGGTGTTGCACTGGTTGATGTAGATGTTGGTCAGCAGGGGATTACG
TGTCCGGGTTCCGTGGCAACAGCTTTTGTGGACAACTATCTCCCCATTGATGAGGGCTTTAACACCGTAATGCCCTTAACCGCCTTCTTTGGCGACAAAACCGTTAATGCCTCTACTCGTGGGCGG
TATCTGGACCTCTGTGCGAGCTTAGCACGTGGCATTATCTCGTTTTCCTTGGCAACTCCGAAATTTGCGGCTGGTGGCGTAATTGTCAACACGATGGGTTGGGTCACCGATATGGGCCTGGATCTGC
TTTTCCAGCTGCTGAGCGTCTTTTCGATTACGCATGTCGTCGTTTGTGGCAGTGGAAATAAGCTGACAGAAACCTTACGCAATGCGGTGATTGACGAGAAAATTATCTTCCTGAAGTATCCGAAACAA
ACCGGGGTCTTTAAACGCAAAGGTAACGTGCGCGATTCTTGGCGTGCGGAACAGATCGTGTCCTACTTCCAAGGGACCAAACGTACCCCTTTACTGTCATATCGTGCCGTGTGCTACGTTAAGGAT
GTACACTTTATCCACGCCTTGAAACTGGAACCGTTAAGCTGGAAAGATGTGGAACCGCTTTCGCTGGCCGCTGTGAGTTGGACCGATAGCCTGGAAGCGGTGAATGATATTAACGTTGCAGGCTTT
ATCGTGTTACTCGAAGTCGGTGAAACCTTCTTTTCGTTTCTGAGTCCAGTTGCGGGTACTCTGCCTAAACCGTTCATTCTTGTATCTCCGACCATTCGCCTTCCTCGCGATAAAGTCCCGCCATTGCA
GGCCCCGCTCGAGCATCACCATCACCATCAC

*Tb*-Clp1-t2-Hisx6 (Amino acid sequence)

MSSNCRTEEFNLERKELKIRFRSSGYVVLTDGAATIFGAPLKKNTRYDFSMCSIPVVSPVACRLHVGGDFTSVVTYIRTDVYDIHAVLDFARHEASKRGVPSLNDTNPGGTSSNSLKEGPWGPRVLVVGD
VNTGKSSLCRSLANMAVASQVHGVALVDVDVGQQGITCPGSVATAFVDNYLPIDEGFNTVMPLTAFFGDKTVNASTRGRYLDLCASLARGIISFSLATPKFAAGGVIVNTMGWVTDMGLDLLFQLLSVFSIT
HVVVCGSGNKLTETLRNAVIDEKIIFLKYPKQTGVFKRKGNVRDSWRAEQIVSYFQGTKRTPLLSYRAVCYVKDVHFIHALKLEPLSWKDVEPLSLAAVSWTDSLEAVNDINVAGFIVLLEVGETFFSFLSPV
AGTLPKPFILVSPTIRLPRDKVPPLQAPLEHHHHHH

*Tb*-Clp1-t3-His Tag (Optimized nucleotide sequence)

ATGGCGAATACCGCGCTGCTGGAGCGCGAATATGCGCTGCCGCCGATGGGTGAACTGTGCTTGGCCACGGCCTCTAGCGGTGGCGCGGCTACCGTGACGTTACTGGCTCAAGTTGAGGACGGC
GGGGAAGAACCTCGCGCAGAGATTTTCGGCACGGAACTCTCCACCGGTGTGGTGGTTCATCTTCCTGTCGCTCGCTCGCTGGCGGTGTTCTCCCCAACAGGTTGTCGCTTAGTGCTCACCGCGAG
TTCAGCCGTTCACCAGATCTGCTATGGTACCACATGCAATGCCACGCGGGCACGCTCGGTAGCGGACATTCACACGCATCTGGAAGTTCAGCGTGTGAAAGCGCGTCGTACCGGCGCAGATGGT
ATTGGCCCGCATGTGCTGTTTGTCGCGGAACGTCGCGCAGTGGGTACAAGCACCTACGTACGCACTCTGATCAACTATGCGGTTCGCCTGGGGTACCATCCGCTGCTTTTGGATGCCTCCGTTGA
AGCGCCACGTTTTGGCTATCCTGGCGTCGTGTCCTTGTATGCAATGCAGTACACCATTGACATTGAGAACGAAATGGCGTTTGTCCCGGGATTACACAGCCATCAAGGGACGAAGAAACATGAAGA
TCCAGCGTTATTCCTGCACATCCTTCGTCAGATGATGCGGCTGTCAACCGAACGCATGGCTCGCTCGGATCGCTGCCGCGTTGGCGGCATTTTCGTAGATTATGGCACCATTAGTCGTGCAGTGGT
CGAGGACGCTGAAGCCTGGGAATGTGCGGAAGAGAAACCCGAAGGTCGCCCGAAAGTTAACCCGCTGGACGTGTTGGTGTCTACGATTTTGGCGGCCGGCATTGACCATGTGTTTGTGGTTGGTA
GTTCATGGCTGCGCTTCAAGATTGCGCAGCGTTTACACCAGGAGTCAGGCGCACAGTCTGAAATCCCGCAAATTACTCCGAGTACCATCACCTGTTCTAACGGCTTGAAAGTTCAGTTGTTCCTGTT
AGATAGCACCGAATGTGGTGCAGTTCCGGATGATGCCTTCTTCAACCGCCAATGCTGGCTGCAGTACTTCTTTGGTTCGCGTACTATGGCTGTCAAACCAACCCTCTTTACCGTGGATGCCAGCCT
TATTCGGCTGGTCACTATTGGGCGTGGTGATACGTCCGGAACAAGCACGTTTATGCCCATGATCGACGATGATTCTGACCACCAAGATCCCACGGTAAGCGGACAGGCTGATGTAGCCCTGACTTA
CGTCCATCCGCAGGATGTGGATATCAAGAATCGTGTACTGGCCCTGAGCACAGCGACCGAACAGGAAGAACTGCCGGATGGGACTTTACAACGCATCCCTTTTGCAGTGTTTGAAAGCCGTCTGA
AACGTGGGCTTCTGATGGGTTTTGCGCTCGTCGAGTCGGTCACAGCAGGCAGTGTGACCCTGCTGACGAATGCTGCAGGAATCCGCAAAGATATTGGCCTCTGCTTTATCGTTACCGACCAGCAG
CTGATGGCCCAAGCCGACGTAGAGCCACCGACTCTCGAGCATCACCATCACCATCAC

*Tb*-Clp1-t3-Hisx6 (Amino acid sequence)

MANTALLEREYALPPMGELCLATASSGGAATVTLLAQVEDGGEEPRAEIFGTELSTGVVVHLPVARSLAVFSPTGCRLVLTASSAVHQICYGTTCNATRARSVADIHTHLEVQRVKARRTGADGIGPHVLF
VAERRAVGTSTYVRTLINYAVRLGYHPLLLDASVEAPRFGYPGVVSLYAMQYTIDIENEMAFVPGLHSHQGTKKHEDPALFLHILRQMMRLSTERMARSDRCRVGGIFVDYGTISRAVVEDAEAWECAEE
KPEGRPKVNPLDVLVSTILAAGIDHVFVVGSSWLRFKIAQRLHQESGAQSEIPQITPSTITCSNGLKVQLFLLDSTECGAVPDDAFFNRQCWLQYFFGSRTMAVKPTLFTVDASLIRLVTIGRGDTSGTSTFM
PMIDDDSDHQDPTVSGQADVALTYVHPQDVDIKNRVLALSTATEQEELPDGTLQRIPFAVFESRLKRGLLMGFALVESVTAGSVTLLTNAAGIRKDIGLCFIVTDQQLMAQADVEPPTLEHHHHHH

*Li*-Clp1-t2-His Tag (Optimized nucleotide sequence)

ATGCAGCCCCGCAAAACGCATACGAGCCTTCACTTGCGGCAAGAAGCGGTGACCATCCAGTGGAATGCGGGTCAGGAAGGCGGCAGTGTTCTGCTGTTATCAGGCAAGGTCGAGTTATTCCGCA
GTAGCCTGACCCGTAATCTGCGCTATGCGTTTCCAGCGGAAGCGTGCATTGTCCTGGAGGCCTTTGGTGACGCCGTAGTGCAAATTGATGGCGATGCAATCACAGTCCAGACTCCCATTTCAGGTG
CGCTCGATGAAATCCACGCCCTGCTGGATACCGCACGTGTTGACGCTATGCTGGCGATTGACGAACGCTCTAAGAAATCCCTTTCGAGCTCTGAAGAGCTGAAAGACTCTTGGCAAGGGCCACGG
GTCCTCGTTGTTGGTGAGAATCAGTGGGAACGTGAAGTCGTGAGTCGTGCCTTACTGAACCTTGCGGTTCGTCATGGCAGCCCGTATGGCATTTGCTACGTAGACGTGGATGTTGCCATGCCTATG
GTGGGTTGTCCAGGAACCGTGTCAGCCGCTTTCGTGGAGGAACCTGTGACAGCGCCGGAAGATTTCAGCGTCATGATGCCGCTGACCTTCTTTCATGGCGCAGCATCGGTAACGAGCGCCACCC
GCAAACGCTATCTGGATCTGTGCGTTTGTGCAGCGCAAGCGGCAACCTCTTTGGGTTTTGCCAACTCCAAATTTGAAGCAGGCGGGTTTCTGATCCACTCGCTGTCTCCGAGTACGGATATTCAGC
ATGACGTGCTGTCCGACGTGATTAGCATCTTCGCCGTAACACACGTTGTTGTCACCGGGGCAGATTGGGAACTCGAAAAGTTTCTCAACAATGCGGTCGTTGGCCGCACTGTGCACTTCGTGCGTT
TGCCGAAATTAGCCGGTGGTCAAAGCCCGAGCGCTGCCGGTGCTGAACAGCGCCGTCGCGCTCAGCTGGAGCATTACTTCTTCGGTACGCCGCGTACTCCACTGATGCCCGTTCGCGGGGTGG
CACGCATGTCGGAACTTGTACTGTTGCATGCGGAAACGTTTGAACCGCTCTCGTGGCGTGAAGTGCCGGATTTAGGCTTAGCAGCTGTCGTTTGGGCGGATACTGCTGCCAGTGCTGTCGAGGCG
AACGTTGCGGGCTTTGTAGCCTTGCTGGAGGTCGGCAAACAGTTCGTGTCCTTTCTGGCGCCTTCCGGAGGAGAACTGCCGAAACCGTTTCTGGTGGTAAGCCCGTCACTGCATCTTCCGCGCGA
GTTGGTGATGCCACTGCCTGTGACCCTCGAGCATCACCATCACCATCAC

*Li*-Clp1-t2-Hisx6 (Amino acid sequence)

MQPRKTHTSLHLRQEAVTIQWNAGQEGGSVLLLSGKVELFRSSLTRNLRYAFPAEACIVLEAFGDAVVQIDGDAITVQTPISGALDEIHALLDTARVDAMLAIDERSKKSLSSSEELKDSWQGPRVLVVGEN
QWEREVVSRALLNLAVRHGSPYGICYVDVDVAMPMVGCPGTVSAAFVEEPVTAPEDFSVMMPLTFFHGAASVTSATRKRYLDLCVCAAQAATSLGFANSKFEAGGFLIHSLSPSTDIQHDVLSDVISIFAV
THVVVTGADWELEKFLNNAVVGRTVHFVRLPKLAGGQSPSAAGAEQRRRAQLEHYFFGTPRTPLMPVRGVARMSELVLLHAETFEPLSWREVPDLGLAAVVWADTAASAVEANVAGFVALLEVGKQFV
SFLAPSGGELPKPFLVVSPSLHLPRELVMPLPVTLEHHHHHH

**Supplementary Table S7. Oligonucleotide sequences used for the phosphorylation assay**

| Oligonucleotide name | RNA/DNA | Oligonucleotide Sequence | Length (nt) |
|---|---|---|---|
| R20-FAM | RNA | 5' - UAAUACGACUCACUAUAGGG -3' (FAM) | 20 |
| R20-comp | RNA | 5' - CCCUAUAGUGAGUCGUAUUA -3' | 20 |
| D20-FAM | DNA | 5' - TAATACGACTCACTATAGGG -3' (FAM) | 20 |
| D20-comp | DNA | 5' - CCCTATAGTGAGTCGTATTA -3' | 20 |

The sequence of oligonucleotide R20-comp is complementary to that of the R20-FAM oligonucleotide. This is also true for the D20-comp and D20-FAM oligonucleotides. FAM, carboxyfluorescein.

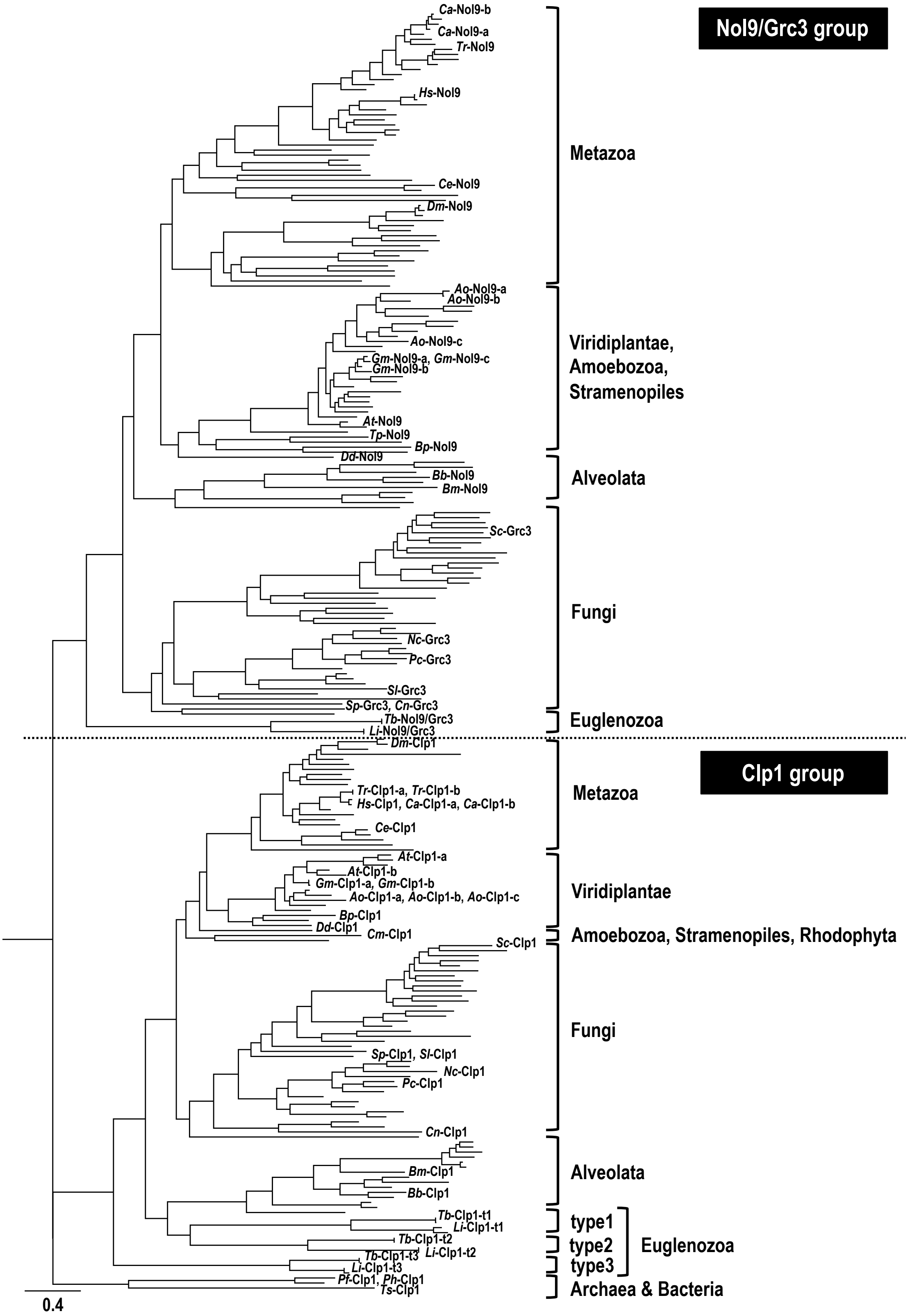


**Supplementary Fig. S1.** Molecular evolutionary phylogenetic tree of Clp1 family proteins (both Clp1 and Nol9/Grc3 groups) examined in this study. A phylogenetic tree based on 254 Clp1 family proteins (both Clp1 and Nol9/Grc3 groups) consisting of 251 protein sequences from 154 eukaryotes, two protein sequences from two archaea, and one protein sequence from a bacterium is shown (see also Supplementary Table S2B). The phylogenetic tree was constructed from their full-length amino acid sequences. Midpoint rooting was applied during tree visualisation. The scale bar under the tree indicates the number of amino acid substitutions per site. The protein names of representative species and taxonomic groups (Kingdom to Phylum) are listed next to the phylogenetic tree. Prokaryota, Archaea, and Bacteria are described as domains. Types 1–3 are protein types of Euglenozoa Clp1 group proteins classified according to their sequence similarities.

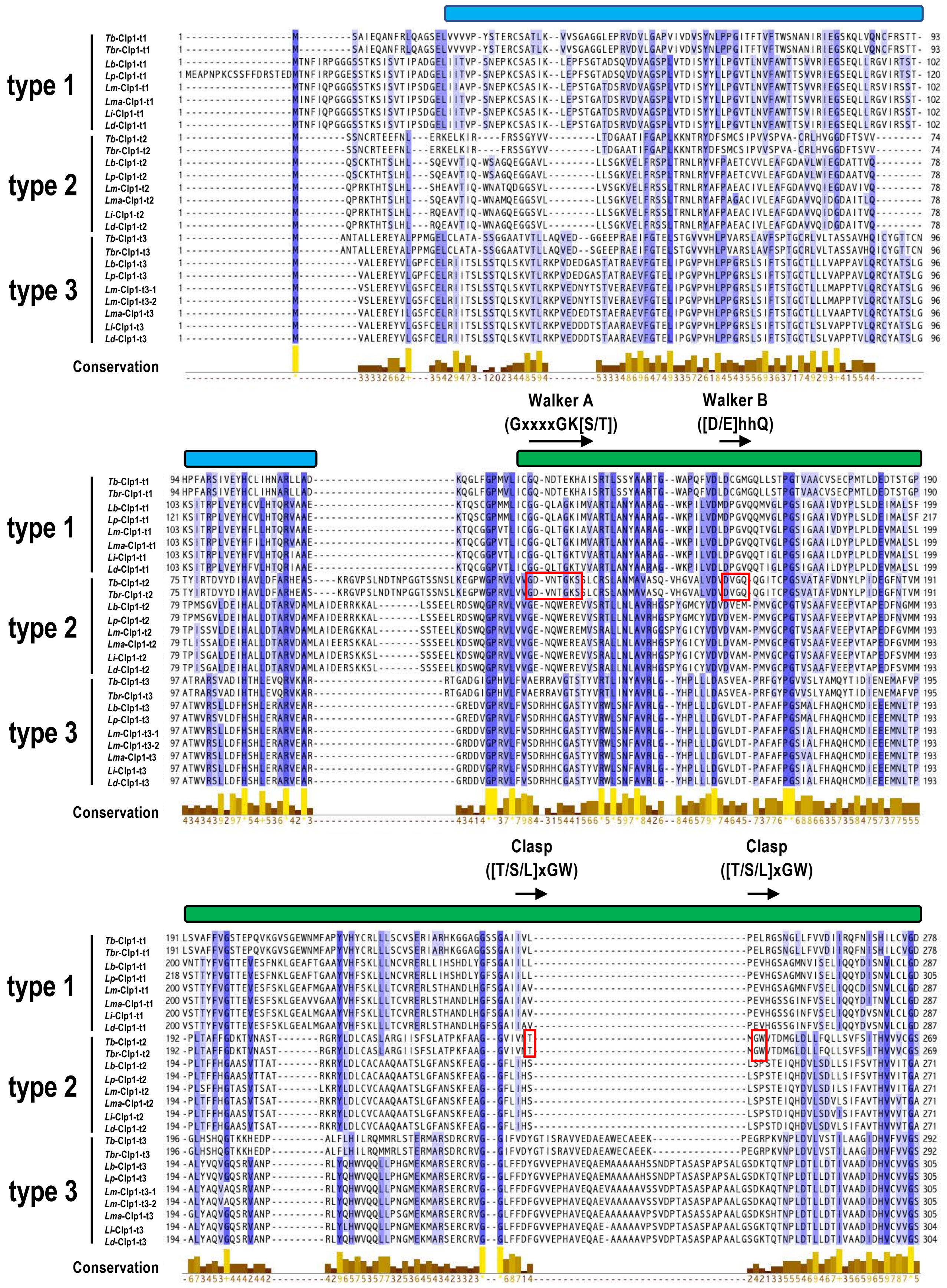


**Supplementary Fig. S2.**

The legend for this figure is placed on the next page.

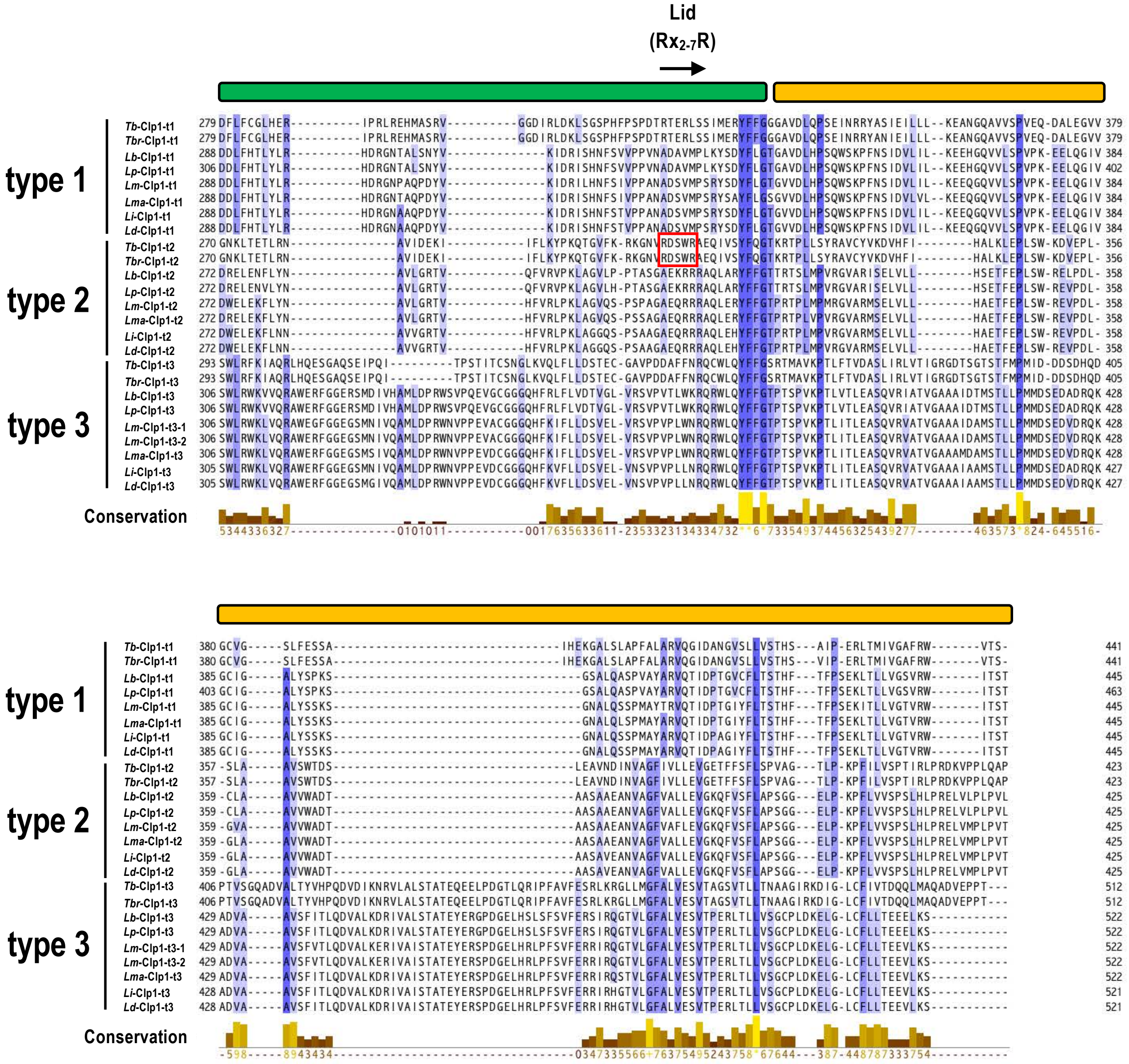


**Supplementary Fig. S2.** Amino acid sequence alignment of Euglenozoa Clp1 group proteins classified as types 1–3. A total of 25 (from eight species; Supplementary Table S3) of 33 Euglenozoa Clp1 family proteins (Supplementary Table S2A), excluding the Nol9/Grc3 group proteins, were used for the sequence alignment. Rectangular boxes above the sequences indicate their protein domain structures (see Supplementary Table S4): Clp1_eN1 (light blue), Clp1_P (green), Clp1_euC1 or Clp1_euC2 or Clp1_euC3 (yellow). The motif name (top), the active site consensus sequence (middle), and its region (arrow at the bottom) are shown for each sequence. The lower part of the figure shows the conservation scores of the alignment. Red boxes indicate areas in which the sequence of the active site is completely conserved. See Supplementary Table S3 for species and protein information used for the analysis.

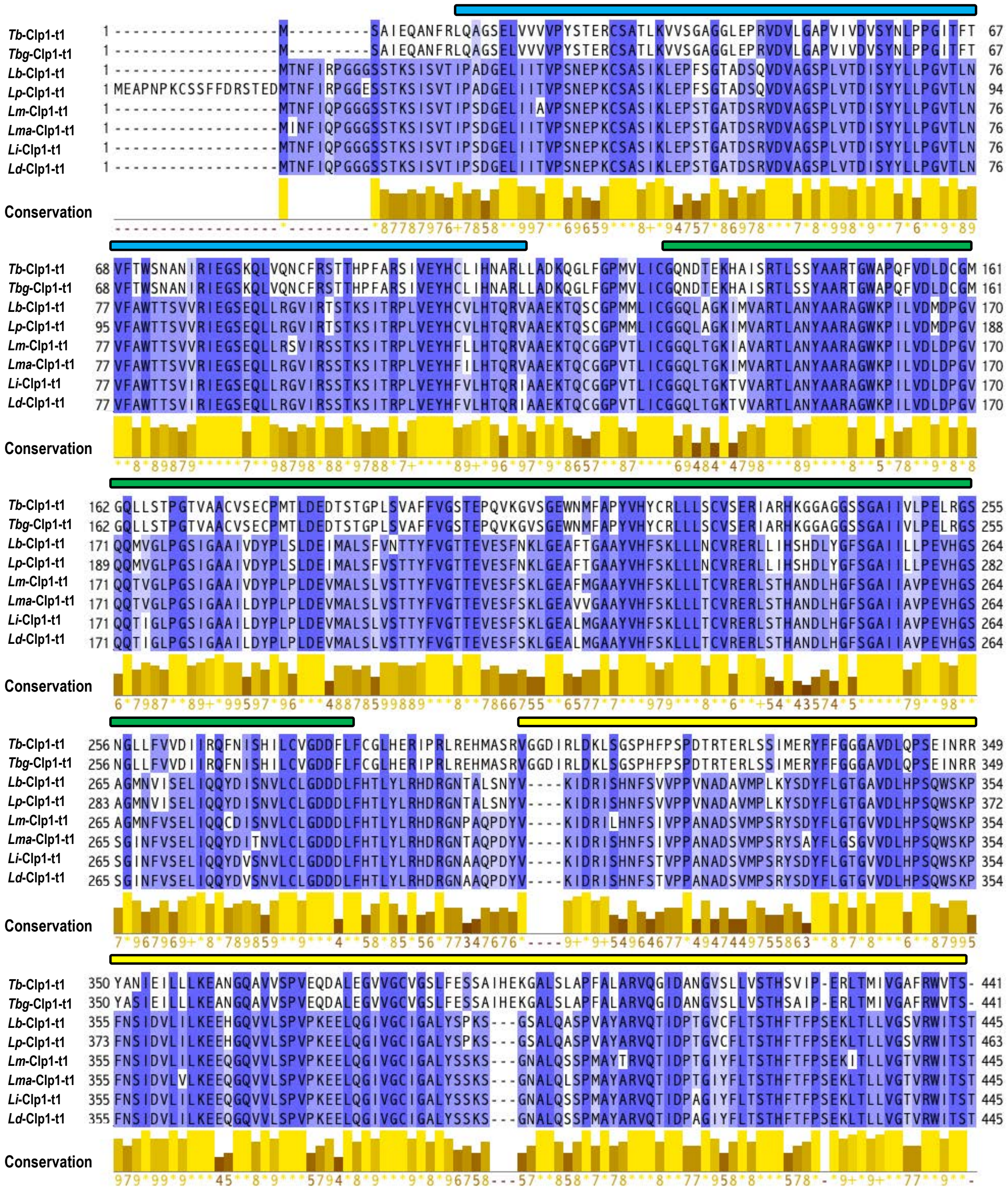

**Supplementary Fig. S3.** Amino acid sequence alignment of the Euglenozoa Clp1 group proteins (type 1). Rectangular boxes above the sequences indicate protein domain structures (Supplementary Table S4): Clp1_eN1 (light blue), Clp1_P (green), and Clp1_euC1 (yellow). The lower part of the figure shows the conservation scores of the alignment. See Supplementary Table S3 for species and protein information used for the analysis.

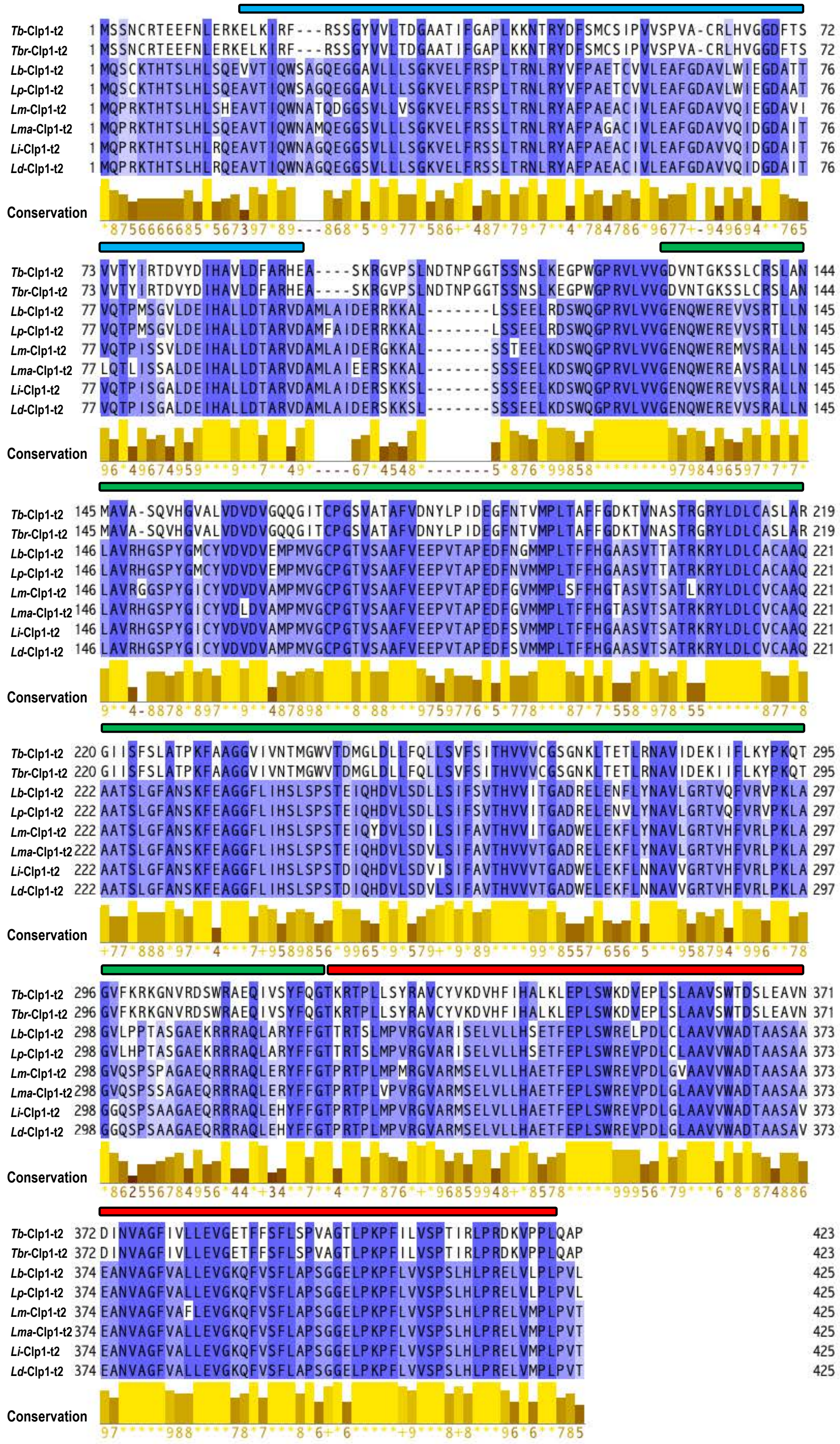

**Supplementary Fig. S4.** Amino acid sequence alignment of the Euglenozoa Clp1 group proteins (type 2). Rectangular boxes above the sequences indicate protein domain structures (Supplementary Table S4): Clp1_eN1 (light blue), Clp1_P (green), and Clp1_euC2 (red). The lower part of the figure shows the conservation scores of the alignment. See Supplementary Table S3 for species and protein information used for the analysis.

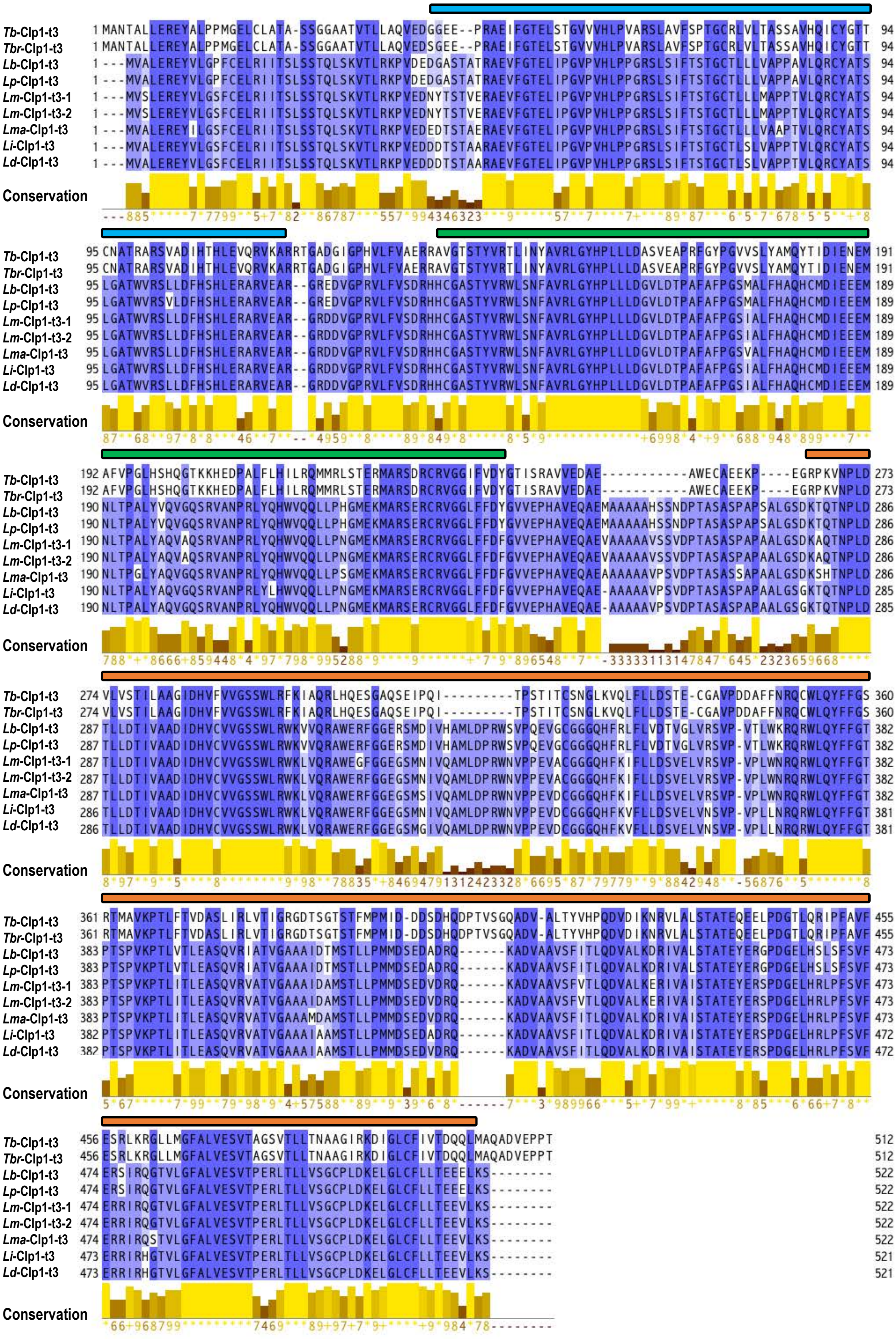


**Supplementary Fig. S5.** Amino acid sequence alignment of the Euglenozoa Clp1 group proteins (type 3). Rectangular boxes above the sequences indicate protein domain structures (Supplementary Table S4): Clp1_eN1 (light blue), Clp1_P (green), and Clp1_euC3 (orange). The lower part of the figure shows the conservation scores of the alignment. See Supplementary Table S3 for species and protein information used for the analysis.

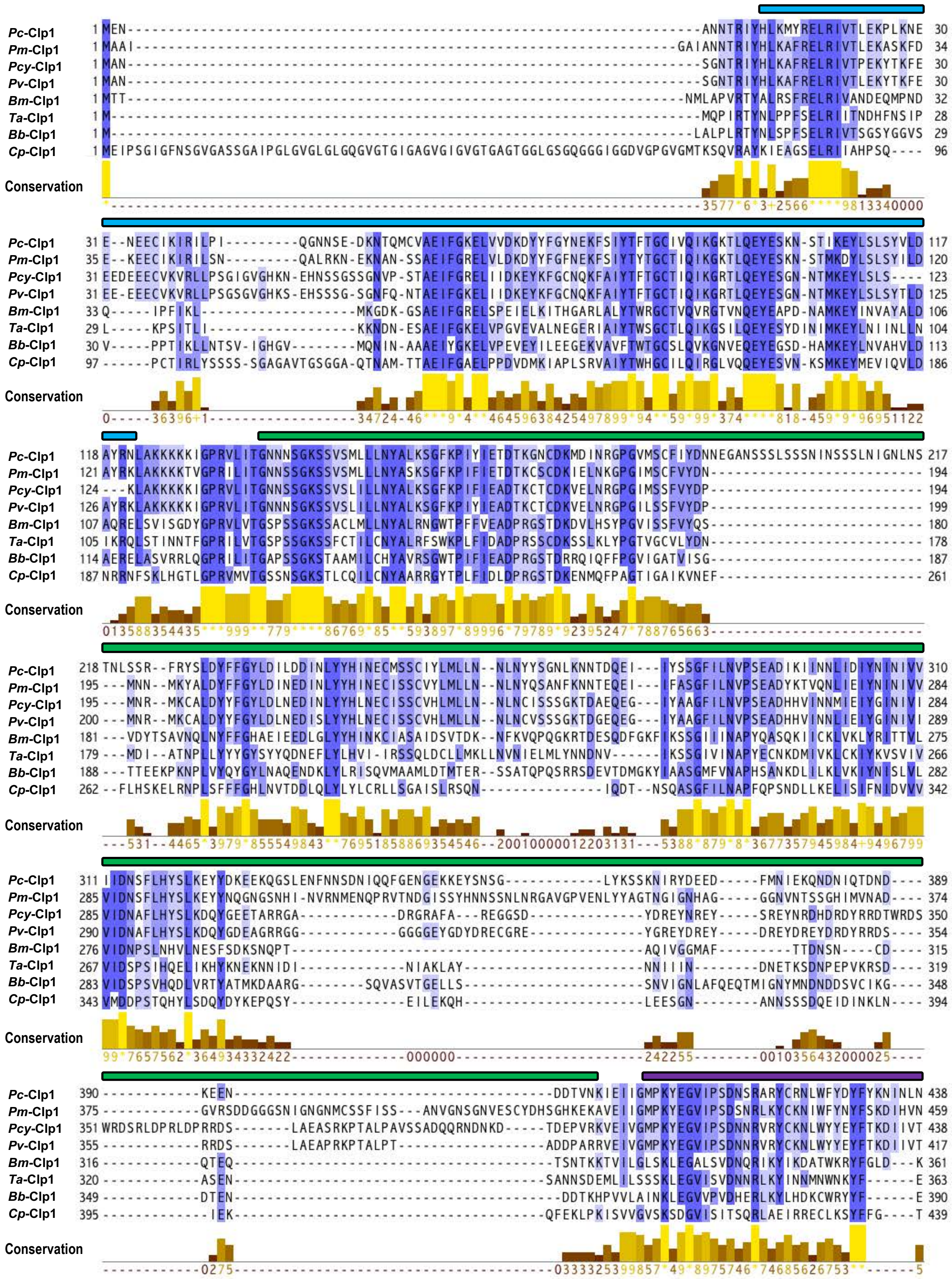


### Supplementary Fig. S6.

The legend for this figure is placed on the next page.

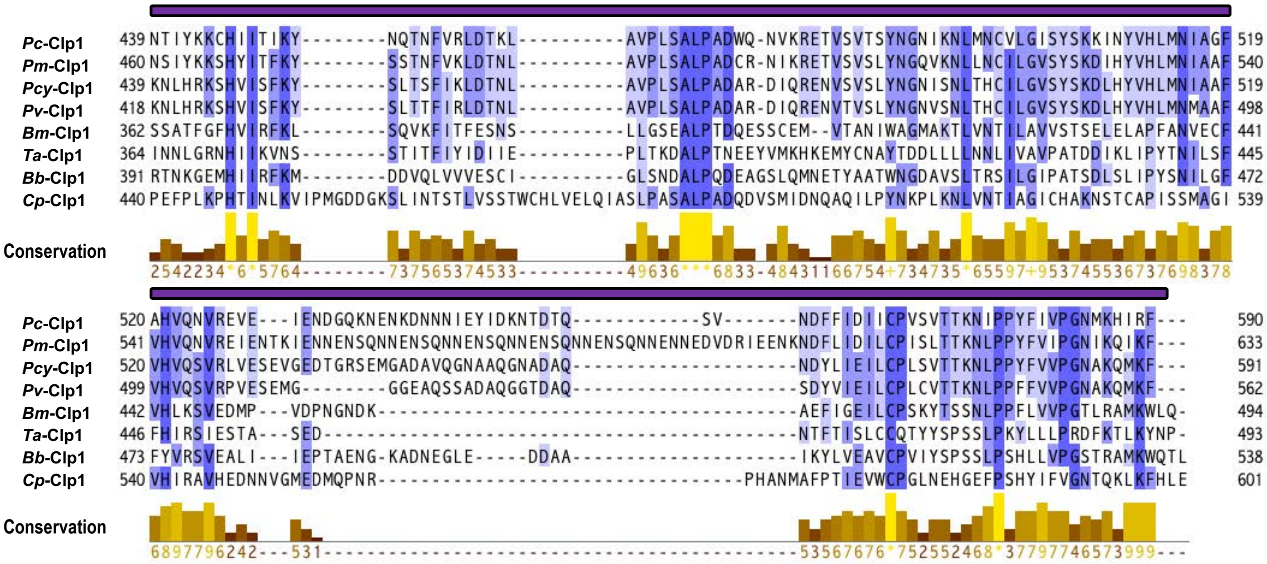

**Supplementary Fig. S6.** Amino acid sequence alignment of the Alveolata Clp1 group proteins. Eight samples of Alveolata Clp1 group proteins (Supplementary Table S3) were randomly selected from 15 samples of Alveolata Clp1 group proteins (Supplementary Table S2B). Rectangular boxes above the sequences indicate protein domain structures (Supplementary Table S4): Clp1_eN1 (light blue), Clp1_P (green), and Clp1_alC (purple). The lower part of the figure shows the conservation scores of the alignment. See Supplementary Table S3 for species and protein information used for the analysis.

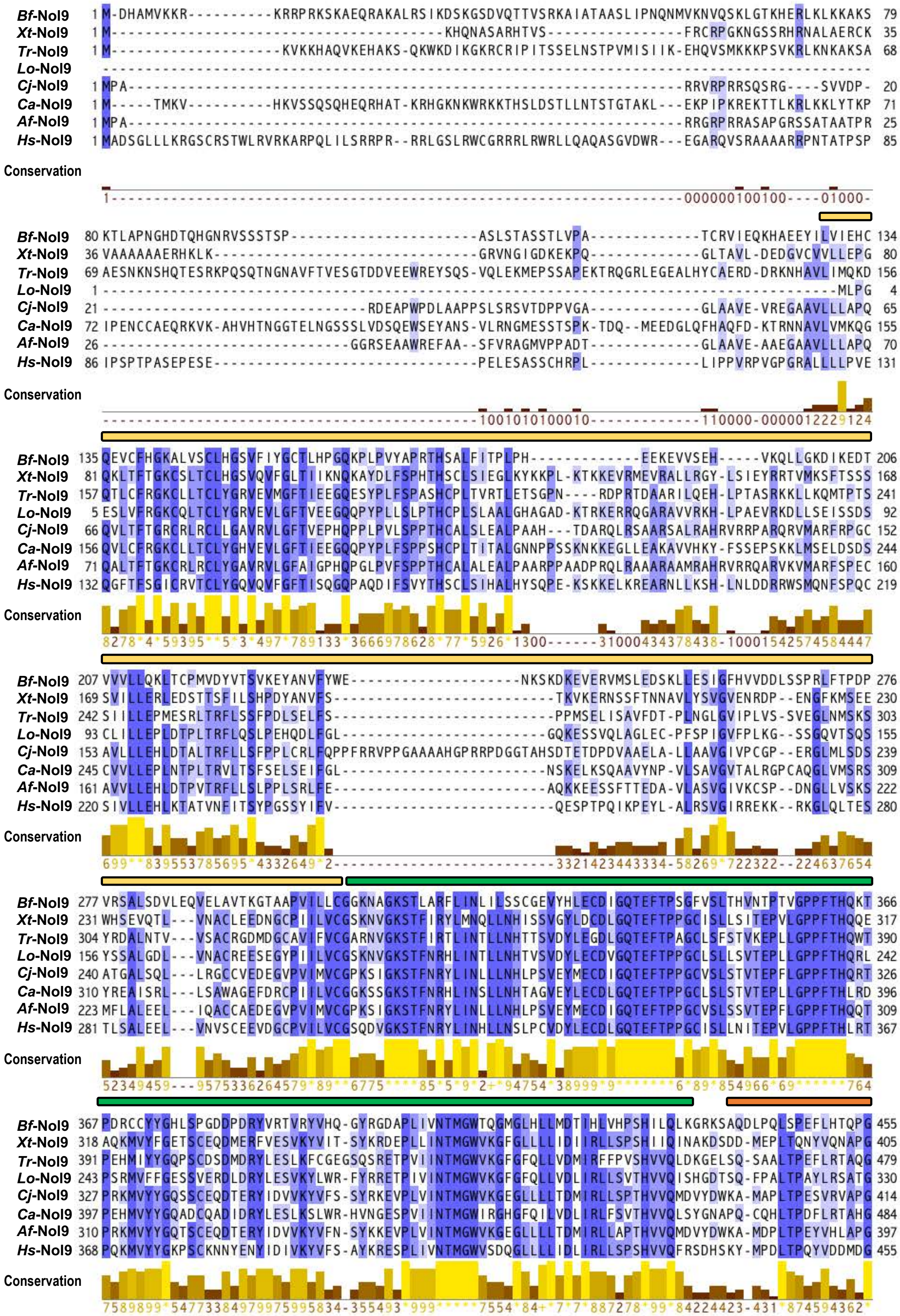

### Supplementary Fig. S7.

The legend for this figure is placed on the next page.

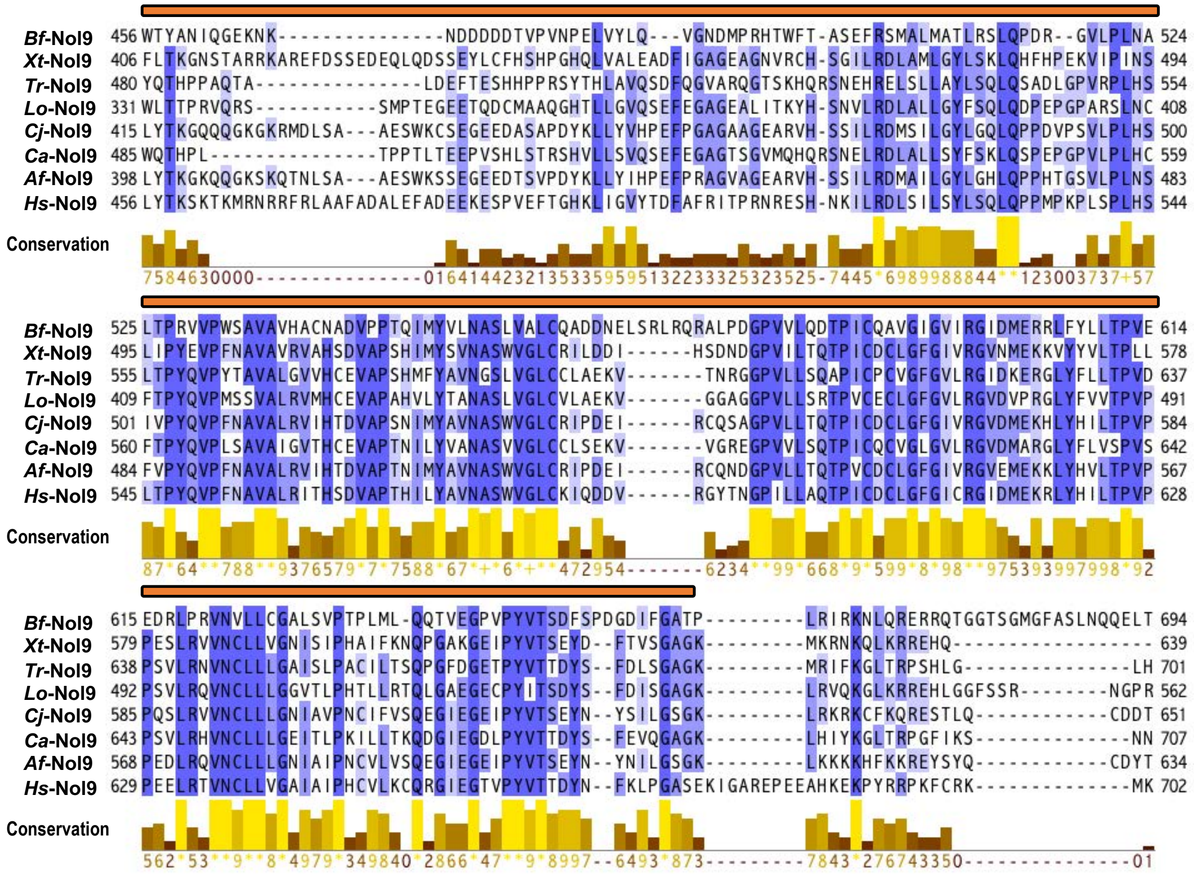

**Supplementary Fig. S7.** Amino acid alignment of the Metazoa Nol9/Grc3 group proteins with the Nol9_eN1 domain. Eight Nol9/Grc3 group proteins (Supplementary Table S3) from 28 Metazoa sequences containing the Nol9_eN1 domain (Figure 3) were used. Rectangular boxes above the sequences indicate protein domain structures (Supplementary Table S4): Nol9_eN1 (light yellow), Clp1_P (green), and Nol9_eC1 (orange). The lower part of the figure shows the conservation scores of the alignment. See Supplementary Table S3 for species and protein information used for the analysis.

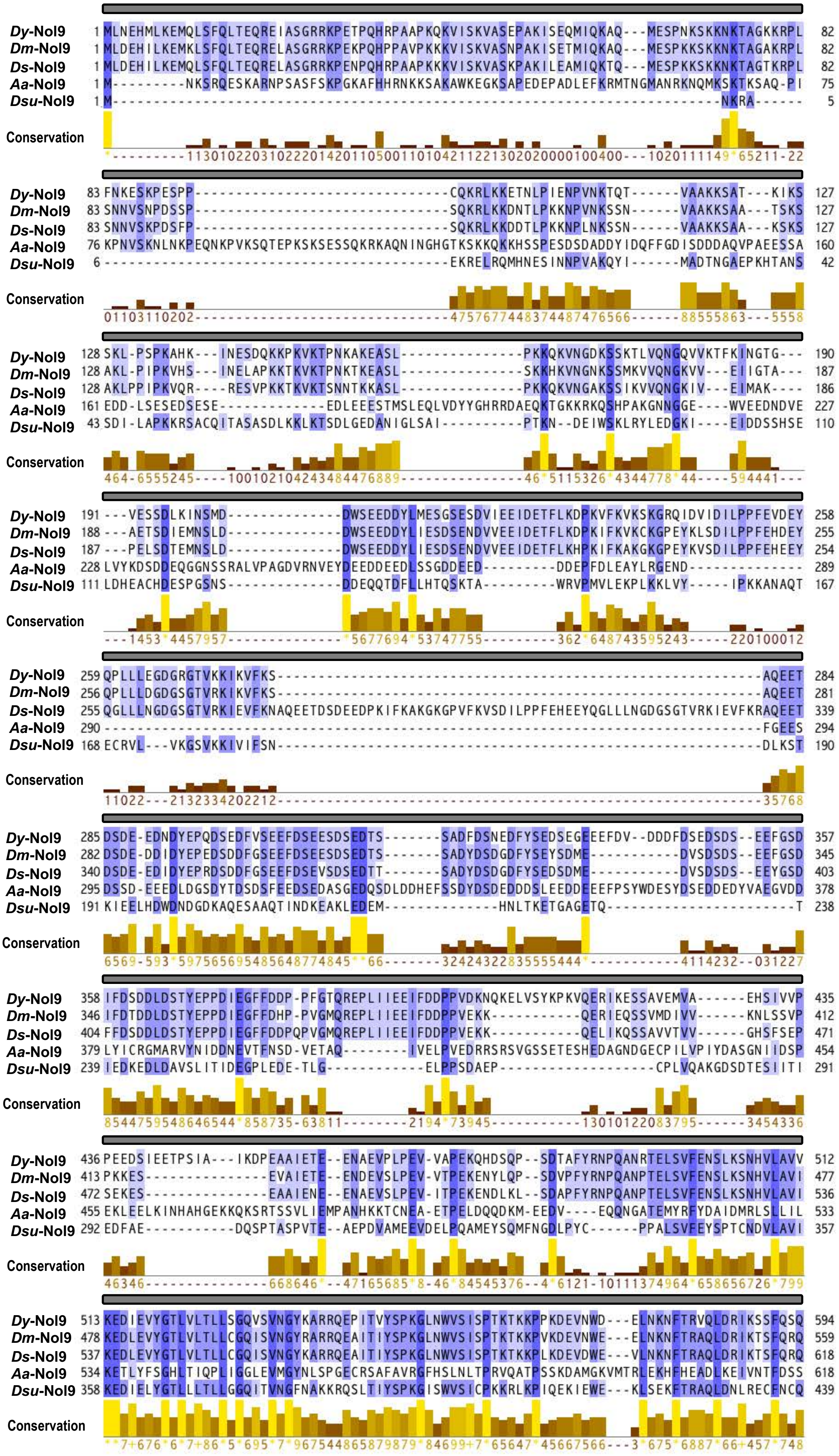


### Supplementary Fig. S8.

The legend for this figure is placed on the next page.

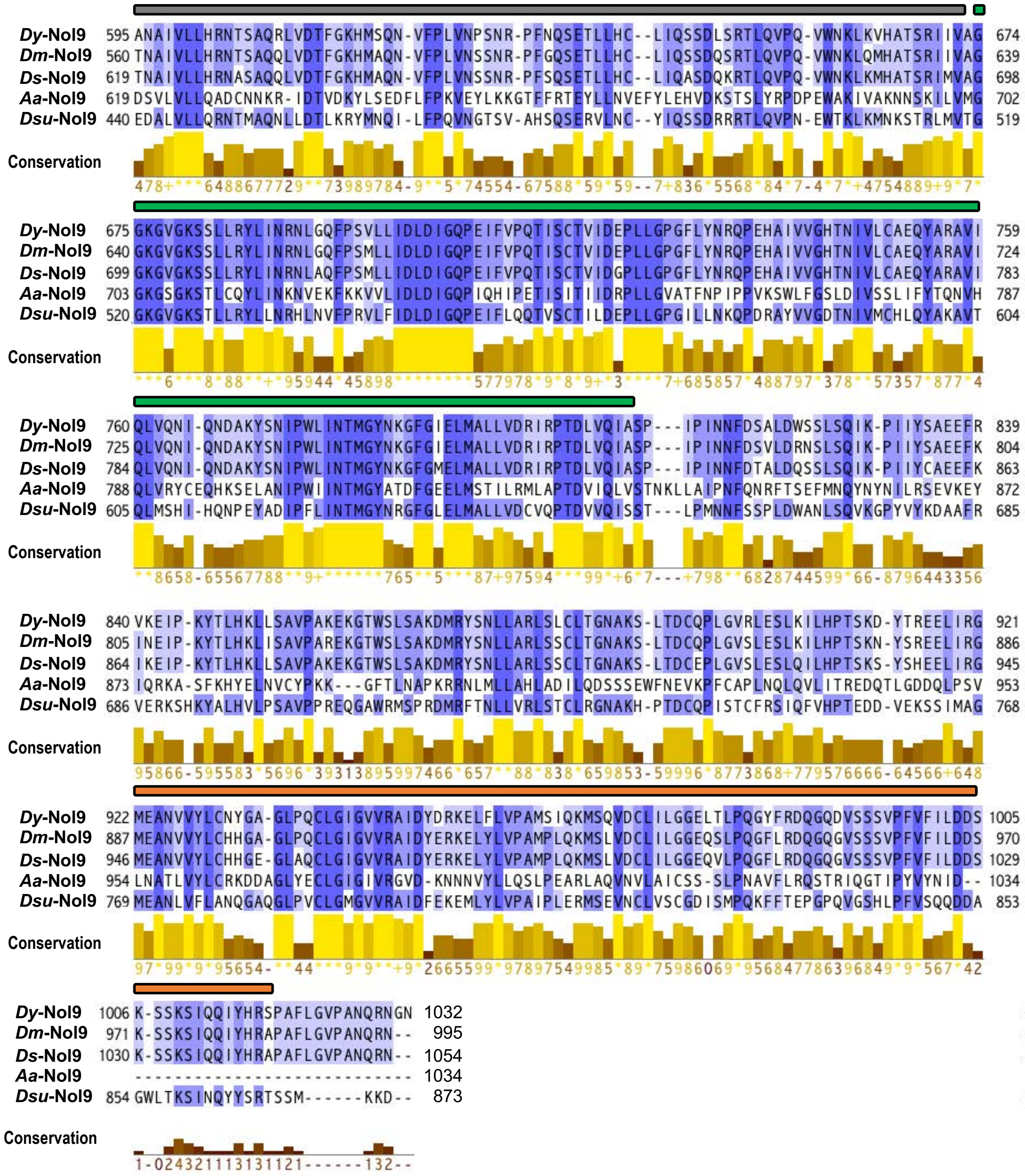

**Supplementary Fig. S8.** Amino acid alignment of the Metazoa Nol9/Grc3 group proteins with the Nol9_eN2 domain. Five Nol9/Grc3 group proteins containing Nol9_eN2 domain were used (Supplementary Table S3). Rectangular boxes above the sequences indicate protein domain structures (Supplementary Table S4): Nol9_eN2 (grey), Clp1_P (green), and Nol9_eC1 (orange). The lower part of the figure shows the conservation scores of the alignment. See Supplementary Table S3 for species and protein information used for the analysis.

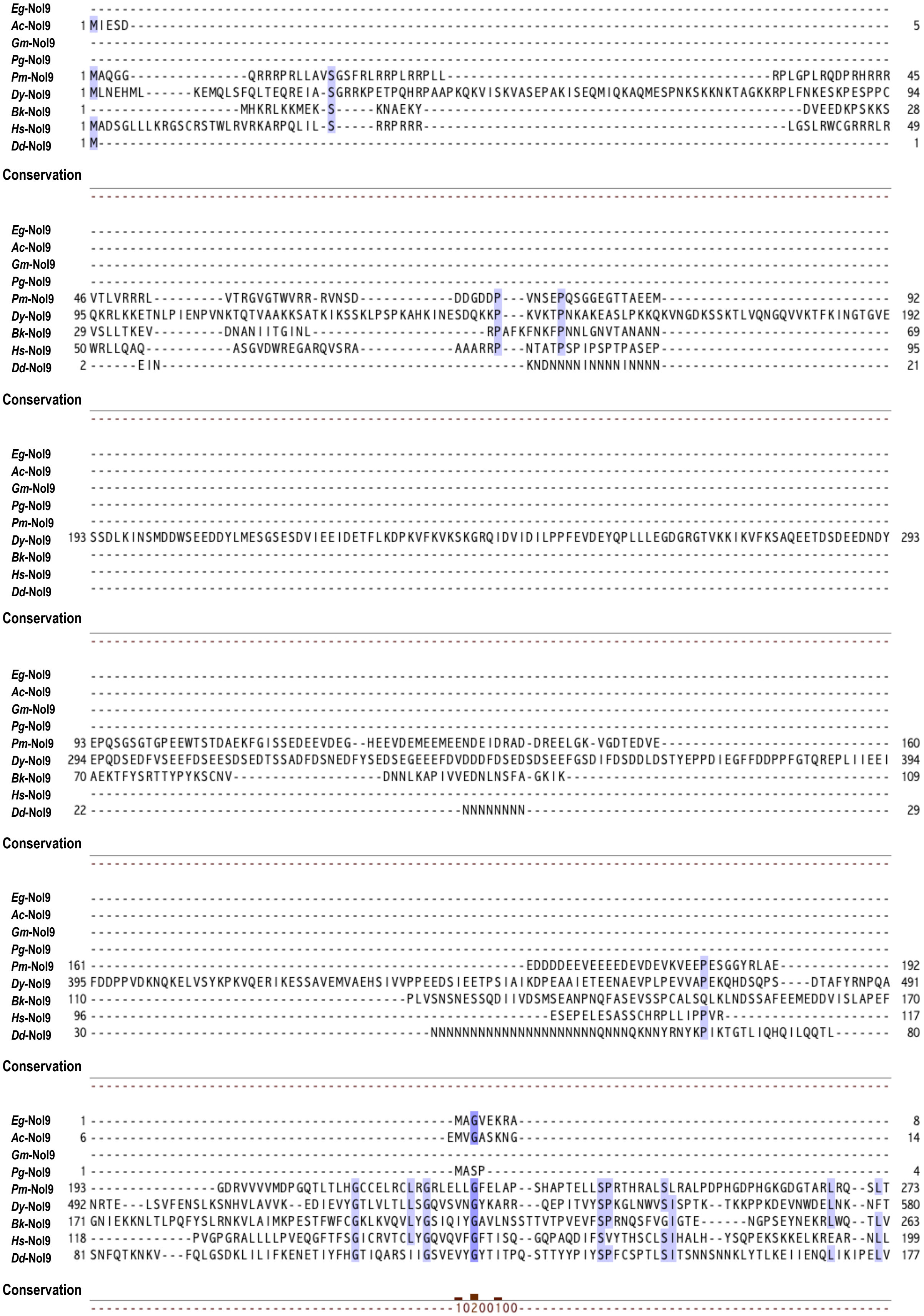


### Supplementary Fig. S9.

The legend for this figure is placed on the next page.

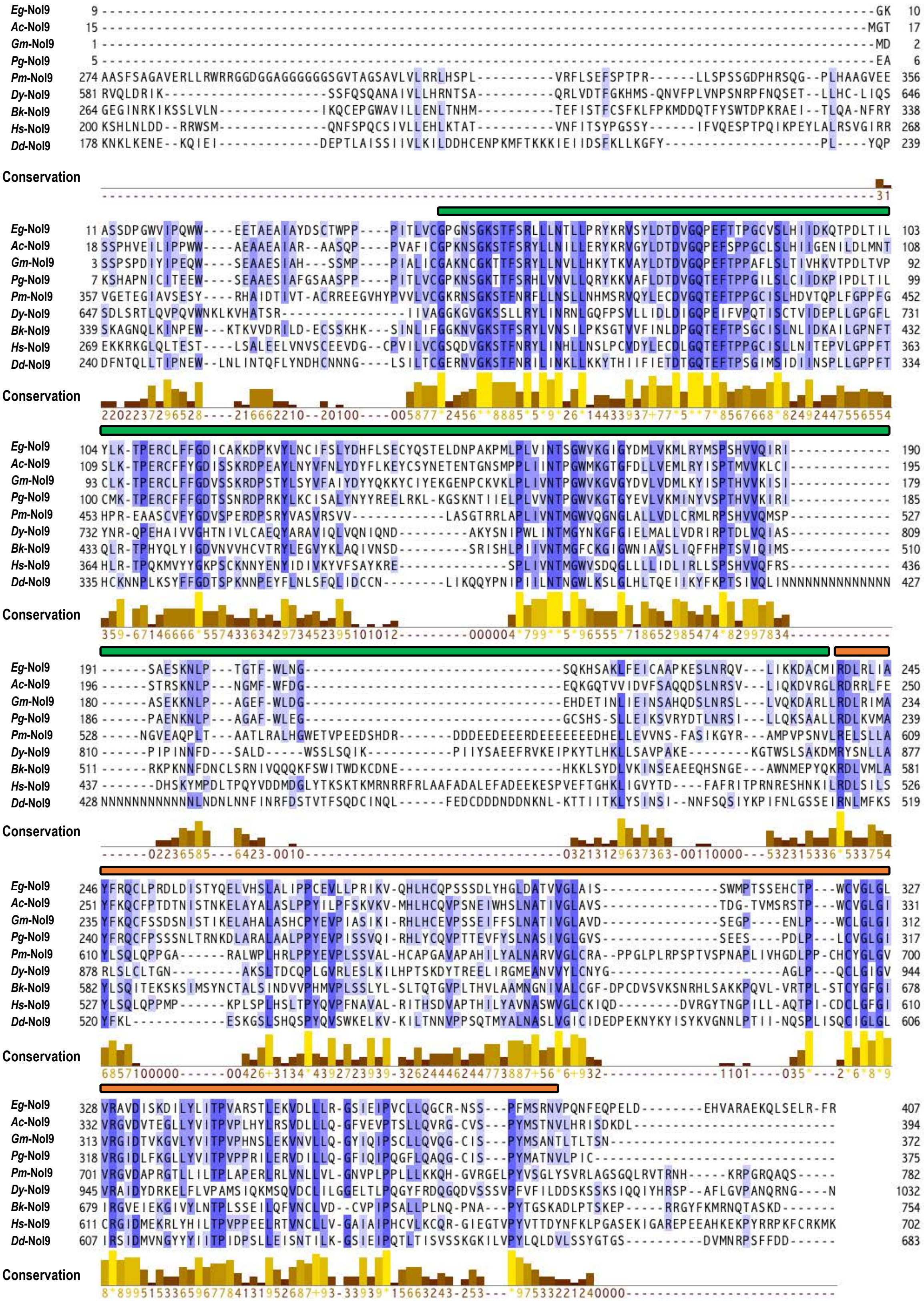

**Supplementary Fig. S9.** Amino acid sequence alignment of Nol9/Grc3 group proteins with the Nol9_eC1 domain from Metazoa, Viridiplantae, and Amoebozoa. Nine examples of Nol9/Grc3 group proteins (Supplementary Table S3) were randomly selected from 55 proteins of Metazoa, 31 proteins of Viridiplantae, and one protein of Amoebozoa (Supplementary Table S2B). Rectangular boxes above the sequences indicate protein domain structures (Supplementary Table S4): Clp1_P (green) and Nol9_eC1 (orange). The lower part of the figure shows the conservation scores of the alignment. See Supplementary Table S3 for species and protein information used for the analysis.

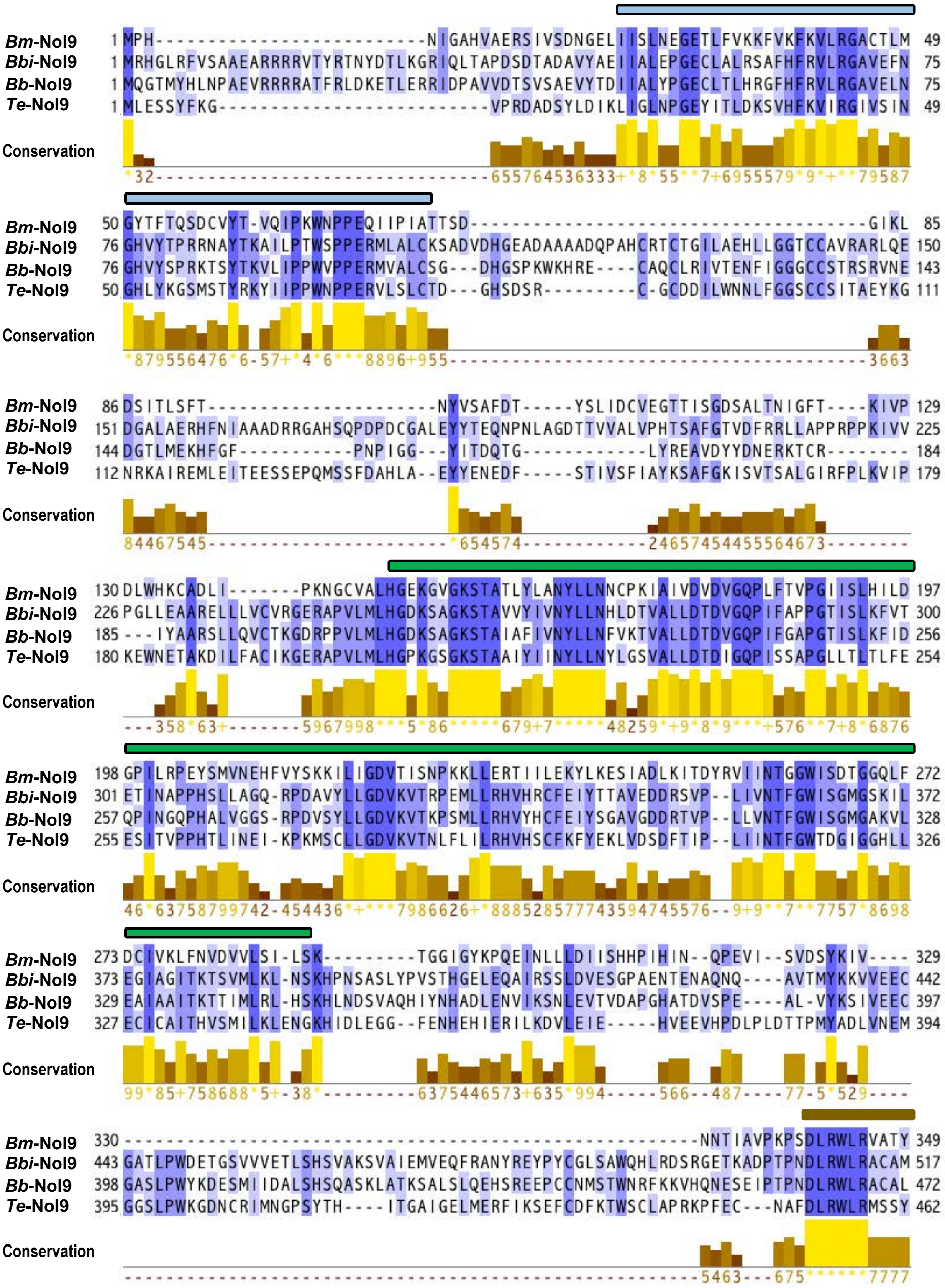


### Supplementary Fig. S10.

The legend for this figure is placed on the next page.

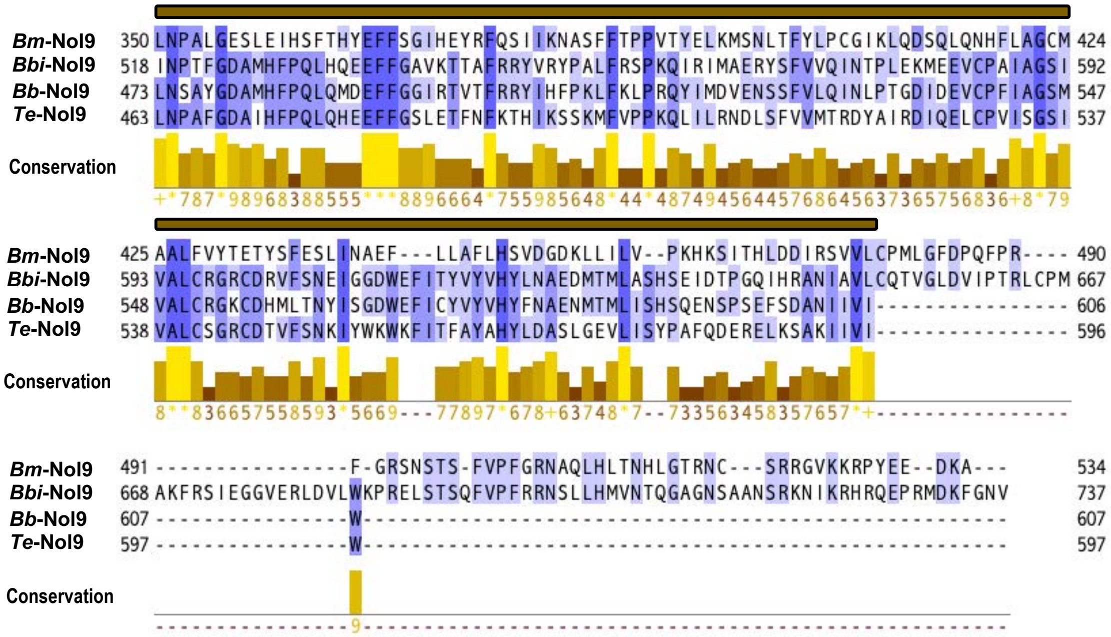

**Supplementary Fig. S10.** Amino acid alignment of Alveolata Nol9/Grc3 group proteins with Nol9_eN3 and Nol9_eC2 domains. Four samples of Alveolata Nol9/Grc3 group proteins (Supplementary Table S3) were used for the alignment. Rectangular boxes above the sequences indicate protein domain structures (Supplementary Table S4): Nol9_eN3 (light blue), Clp1_P (green), and Nol9_eC2 (light brown). The lower part of the figure shows the conservation scores of the alignment. See Supplementary Table S3 for species and protein information used for the analysis.

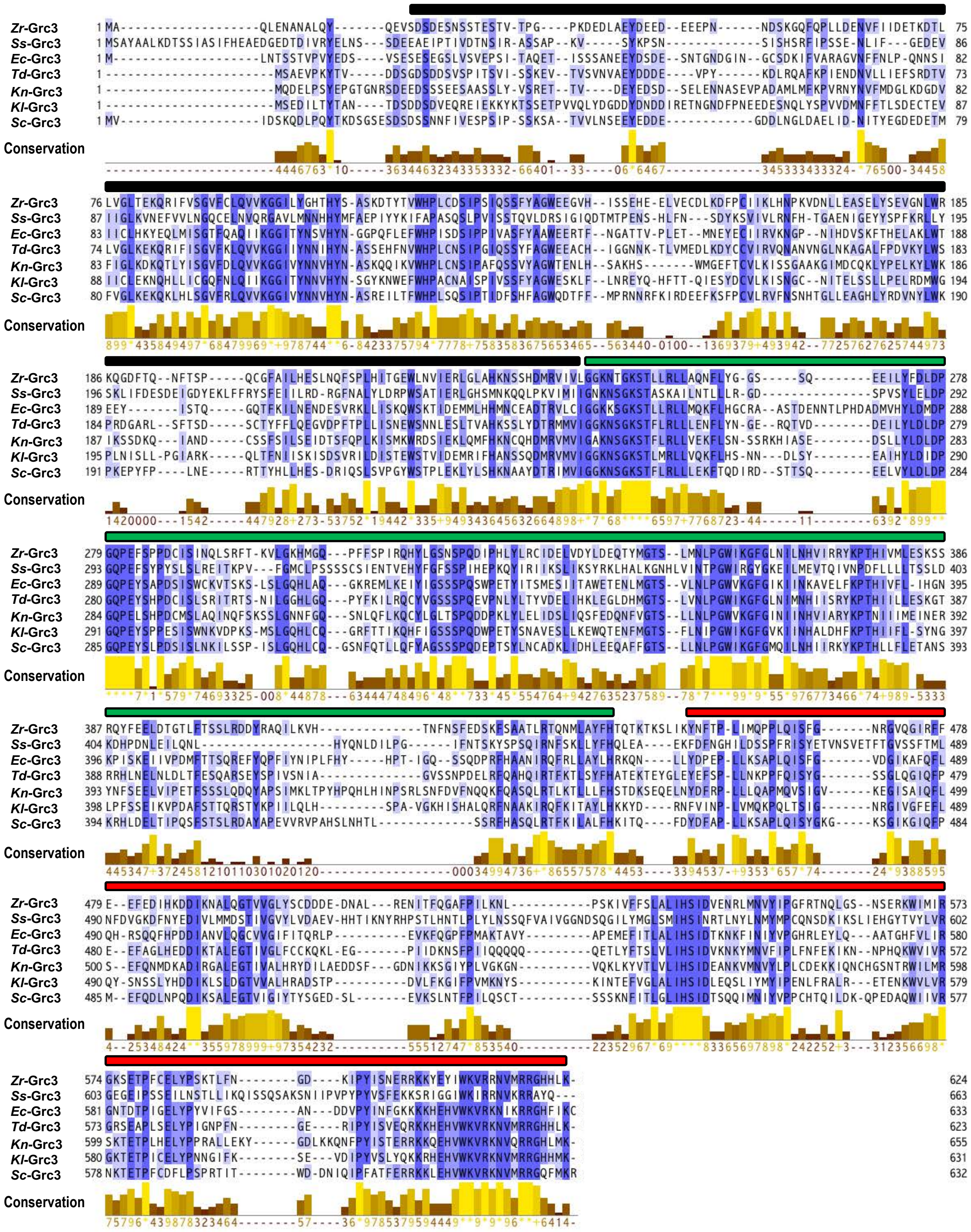


**Supplementary Fig. S11.** Amino acid alignment of the fungal Nol9/Grc3 group proteins with Grc3_eN1 and Grc3_eC1 domains. Seven samples of fungal Nol9/Grc3 group proteins with Grc3_eN1 and Grc3_eC1 domains (Supplementary Table S3) were used for the alignment. Rectangular boxes above the sequences indicate protein domain structures (Supplementary Table S4): Grc3_eN1 (black), Clp1_P (green), and Grc3_eC1 (red). The lower part of the figure shows the conservation scores of the alignment. See Supplementary Table S3 for species and protein information used for the analysis.

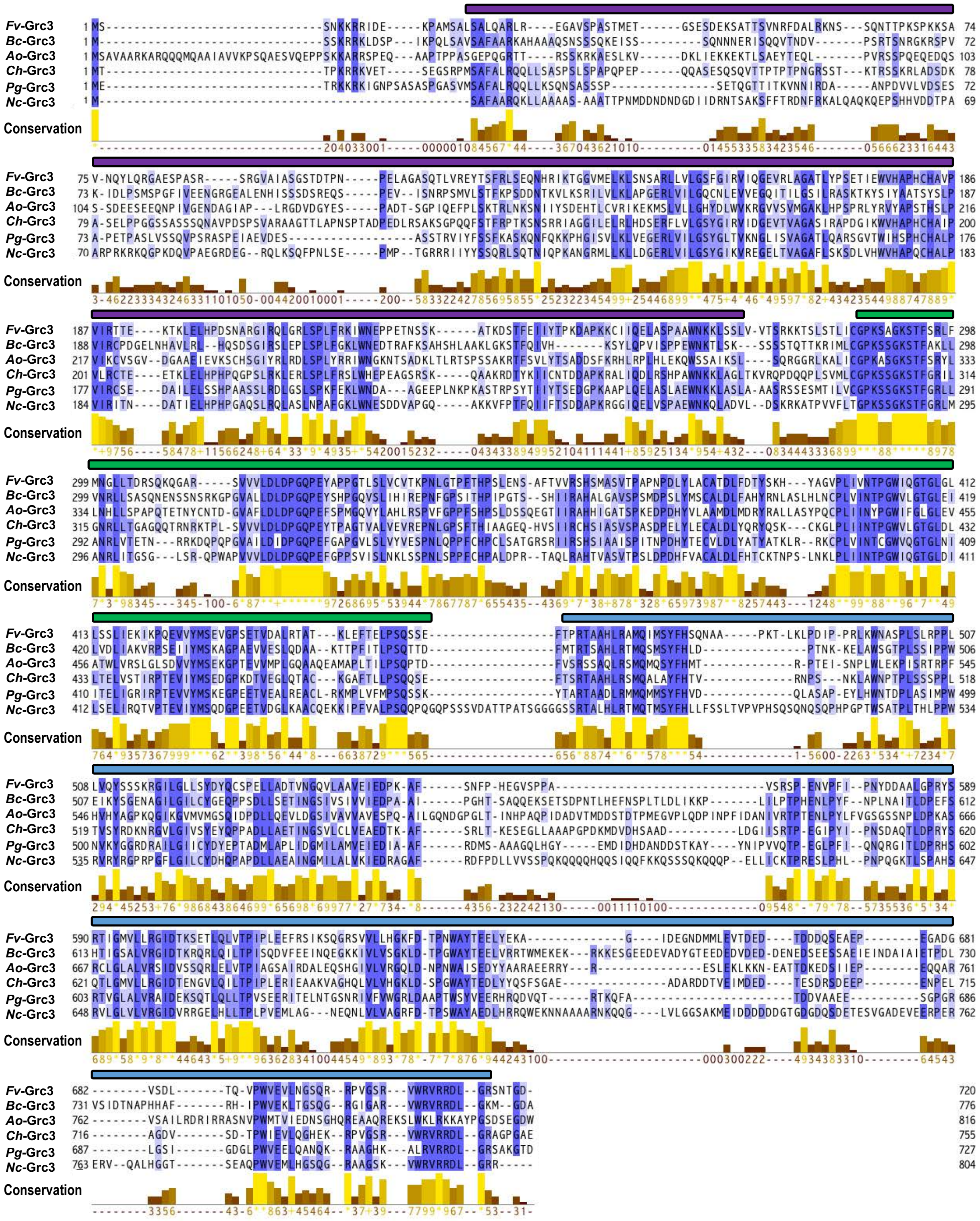


**Supplementary Fig. S12.** Amino acid alignment of fungal Nol9/Grc3 group proteins with Grc3_eN2 and Grc3_eC2 domains. Six examples of fungal Nol9/Grc3 group proteins containing Grc3_eN2 and Grc3_eC2 domains (Supplementary Table S3) were used for the alignment. Rectangular boxes above the sequences indicate protein domain structures (Supplementary Table S4): Grc3_eN2 (purple), Clp1_P (green), and Grc3_eC2 (blue). The lower part of the figure shows the conservation scores of the alignment. See Supplementary Table S3 for species and protein information used for the analysis.

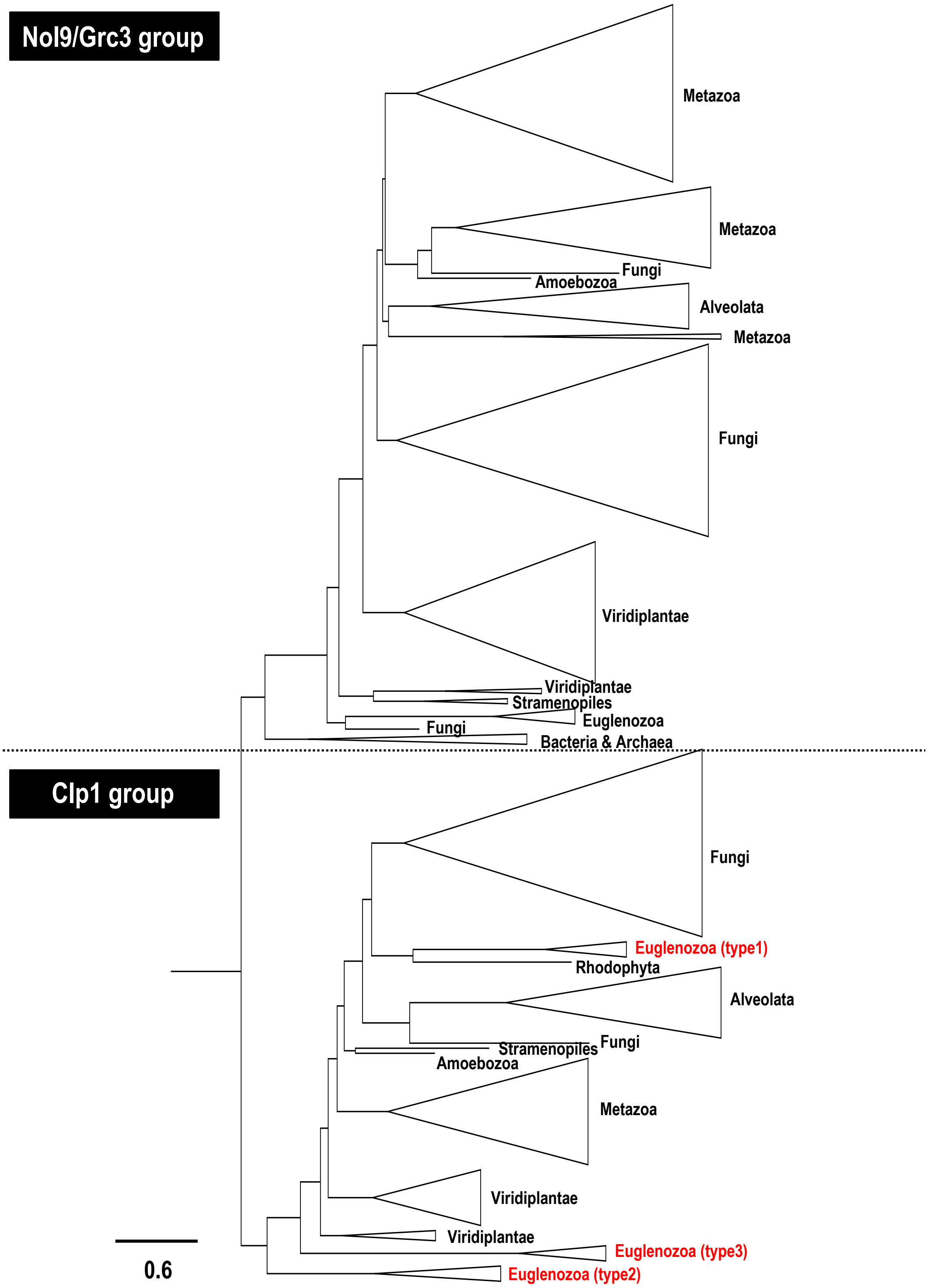


**Supplementary Fig. S13.** Molecular evolutionary phylogenetic tree of Clp1 family proteins (both Clp1 and Nol9/Grc3 groups) constructed from the amino acid sequences of the PNK domain. A molecular evolutionary phylogenetic tree of Clp1 family proteins constructed from the amino acid sequences of the PNK domain, which is commonly conserved in Clp1 family proteins, is shown. Among 254 Clp1 family proteins (Supplementary Figure S2B), 253 Clp1 family proteins from 250 eukaryotes, two archaea, and one bacterium were used, excluding one species, *Nasonia vitripennis* (RefSeqID: XP_008207576.1), with a partially incomplete PNK domain region. Midpoint rooting was applied during tree visualisation. The scale bar under the tree indicates the number of amino acid substitutions per site. The protein names of representative species and taxonomic groups (Kingdom to Phylum) are listed next to the phylogenetic tree. Prokaryota, Archaea, and Bacteria are described as domains. Types 1–3 are protein types of Euglenozoa Clp1 group proteins classified according to their sequence similarity. The same taxa are grouped in triangles, the sizes of which reflect the number of sequences.

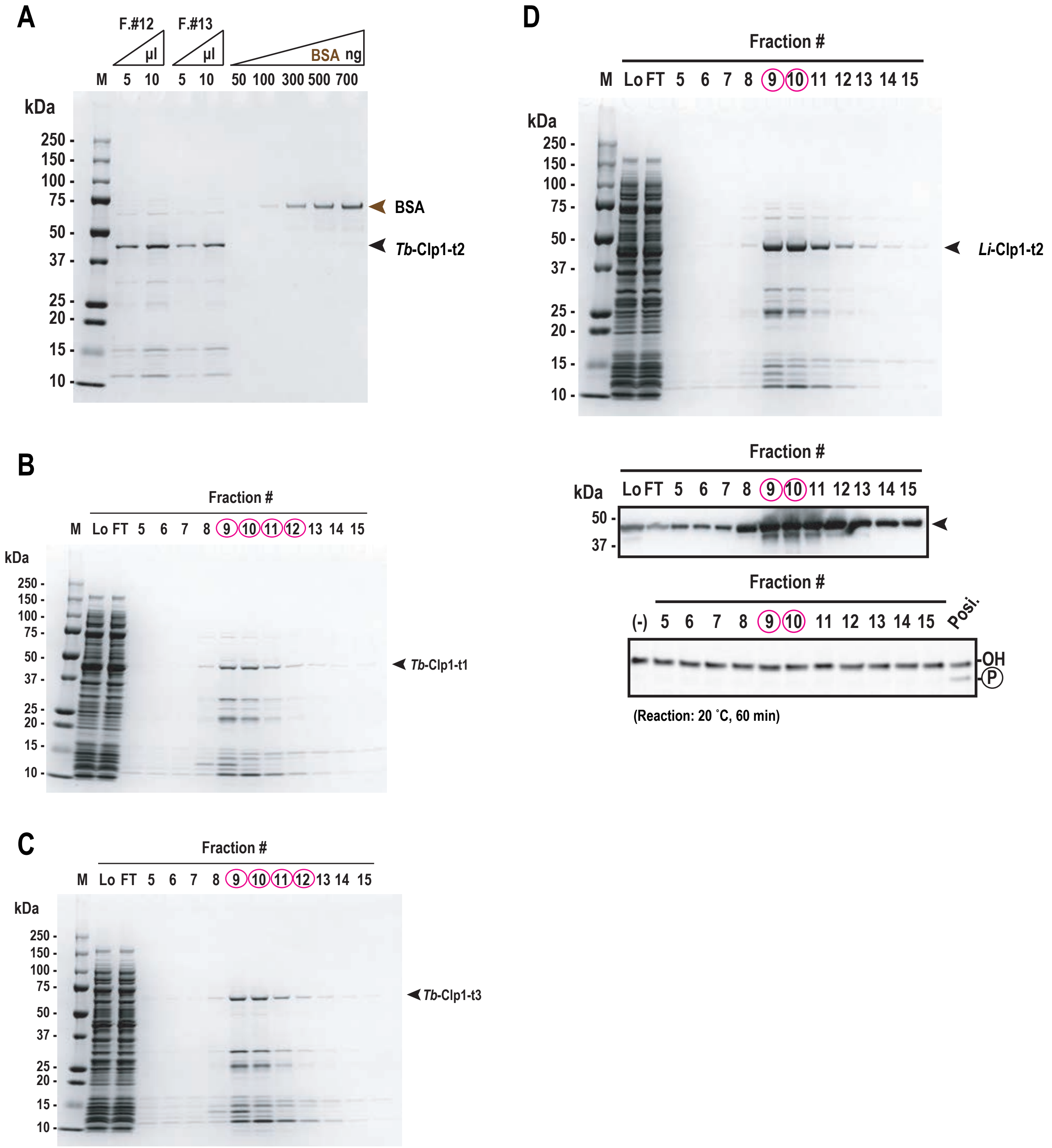


**Supplementary Fig. S14.** Purification of recombinant Euglenozoa Clp1 group proteins and their PNK activities (supporting data). (A) Determination of concentration of partially purified *Tb*-Clp1-t2 protein using bovine serum albumin (BSA) at known concentrations (50–700 ng) as the standard. Samples were subjected to 10%–20% SDS-PAGE and stained with Coomassie Brilliant Blue. F.: fraction. (B) Each fraction of the recombinant *Tb*-Clp1-t1 and (C) *Tb*-Clp1-t3 proteins purified with HisTrap HP affinity chromatography was subjected to 10%–20% SDS-PAGE and stained with Coomassie Brilliant Blue. (D) Each fraction of recombinant *Li*-Clp1-t2 protein purified with HisTrap HP affinity chromatography was subjected to 10%–20% SDS-PAGE and stained with Coomassie Brilliant Blue (top). Western blotting analysis with anti-His-tag antibody (middle), and PNK activity using ssRNA as the substrate (bottom). Recombinant *Tb*-Clp1-t2 protein was used as the positive control. Protein peaks on column chromatography are indicated by red circles. Arrows indicate the position of each protein (BSA, light brown; each recombinant protein. black).
